# SiRCle (Signature Regulatory Clustering) model integration reveals mechanisms of phenotype regulation in renal cancer

**DOI:** 10.1101/2022.07.02.498058

**Authors:** Ariane Mora, Christina Schmidt, Brad Balderson, Christian Frezza, Mikael Bodén

## Abstract

Clear cell renal cell carcinoma (ccRCC) tumours develop and progress via complex remodelling of the kidney epigenome, transcriptome, proteome, and metabolome. Given the subsequent tumour and inter-patient heterogeneity, drug-based treatments report limited success, calling for multi-omics studies to extract regulatory relationships, and ultimately, to develop targeted therapies. However, current methods are unable to extract nonlinear multi-omics perturbations.

Here, we present SiRCle (**Si**gnature **Re**gulatory **Cl**ust**e**ring), a novel method to integrate DNA methylation, RNA-seq and proteomics data. Applying SiRCle to a case study of ccRCC, we disentangle the layer (DNA methylation, transcription and/or translation) where dysregulation first occurs and find the primary biological processes altered. Next, we detect regulatory differences between patient subsets by using a variational autoencoder to integrate omics’ data followed by statistical comparisons on the integrated space. In ccRCC patients, SiRCle allows to identify metabolic enzymes and cell-type-specific markers associated with survival along with the likely molecular driver behind the gene’s perturbations.

## Introduction

Clear Cell Renal Cell Carcinoma (ccRCC) is the most prevalent form of kidney cancer and accounts for 70% of renal malignancies^1^. It is now established that most ccRCC cases are driven by the loss of the Von Hippel-Lindau (VHL) tumour suppressor gene, in turn leading to the activation of the transcription factor (TF) Hypoxia-Inducible Factors (HIFs)^2^. Although the activation of the HIF pathway is considered to be a driving event in VHL mutant ccRCCs, VHL mutations alone are insufficient for ccRCC formation and cooperating mutations such as PBRM1, BAP1, KDM5C and SETD2 are necessary for full transformation^3^. Hence, ccRCC transformation and progression have been reported to affect multiple regulatory layers, beyond transcription, including the epigenome, via changes in DNA methylation of CpG islands^4^, the proteome, via alterations in the mammalian target of rapamycin (mTOR) pathway^3^ that in turn affects protein synthesis^5^, and the metabolome, whose dysregulation is considered a hallmark of ccRCC^6–8^.

Due to the complexity of gene regulation in ccRCC, molecular analyses for patient stratification and drug treatment based exclusively on genomic or transcriptomics analysis have been unsuccessful^9^. For example, protein expression via (post-) translational regulation is altered in cancer^10^ and there is a lack of correlation between transcriptome and proteome^11^. Hence, it is difficult to predict pathway activity such as TFs driving disease states from transcriptomics data alone, suggesting that data from distinct regulatory layers need to be analysed jointly. Furthermore, patient’s responses to different RCC treatments can be dependent on the mutational and methylation landscape of the patients^12^. To overcome these issues, consortia such as CPTAC (Clinical Proteomic Tumor Analysis Consortium) are leading efforts to produce multi-omics datasets with various patient demographics^13^. Clark *et. al*. reported the first extensive dataset of ccRCC patients that included tumour and normal RNA-seq and proteomics data, and tumour DNA methylation data^14^. Using largely correlative analyses, Clark *et. al* proposed a stratification of ccRCC patients for personalised therapeutic interventions based on the differential RNA and protein profiles of the patients, along with the copy number variation and phosphoproteome^14^. However, it remains unclear if in ccRCC a gene of interest is regulated at the level of DNA-methylation, transcription or translation, and at what regulatory level (henceforth “layer”) dictates the cellular phenotype of ccRCC. Addressing this question is vital to identify determinants that explain disease states and to subsequently develop effective anticancer therapies.

Data integration has become a core component of multi-omic analyses but remains a challenge owing to dataset heterogeneity, noise, and non-linearities between genetic and epigenetic interactions^15^. Many approaches exist to integrate bulk multi-omic assays to understand cancer genomics; they are typically based on standard statistical methods, clustering, or linear matrix factorisations to integrate different biological data^16,17^. Recently, deep learning methods, such as Variational Autoencoders (VAEs)^18^, have been applied to detect non-linear patterns across patients in single genomic data types in cancer patients^19,20^. VAEs have been used for differential analysis, cancer subtype identification, data integration^21,22^, and survival analysis^23^, involving both bulk^19^ and single cell^24^ data, and on a variety of assays (ATAC^21^, mRNA^19,25,26^, DNA methylation^26^).

VAEs present a machine learning approach to extract latent features that summarise variation across the dataset^27^. Recent research suggests that latent embeddings can be used to identify co-regulated gene groups^28^; however, such analyses have yet to be adapted for application on cancer or patient data. Other integration methods enable biological signals to be extracted along latent embeddings, typically in the forms of correlative or linear relationships^29–34^, statistics on these bulk-cancer embeddings are not often performed. Yet, this is becoming increasingly important as it has been shown that patient’s demographics such as age^35–37^, gender^38,39^ and ethnicity^40^ alter the gene expression profile in ccRCC as in many other tumour types. Furthermore, it is important to stratify patients on stage^41^, and mutational profile^42^ since these factors are known to affect survival. To date, studies involving GWAS data suggest principal component analysis (PCA) embedding statistics enable complex relationships to be identified, opening an avenue for multi-omic patient changes to be extracted^43,44^.

Here we present “**Si**gnature **R**egulatory **Cl**ust**e**ring’’ (SiRCle), a method for integrating DNA methylation, mRNA, and protein data at the gene level to deconvolute the association between dysregulation within and across possible regulatory layers (DNA methylation, transcription and/or translation), which is available as a Python package. Using SiRCle, we disentangled the regulatory layer behind metabolic rewiring in ccRCC and showed that glycolysis is regulated by DNA hypomethylation, whilst mitochondrial enzymes and respiratory chain complexes are translationally suppressed. Moreover, we identified that HIF1A was likely driving expression changes in glycolytic enzymes. We also explored the heterogeneity between ccRCC patient subpopulations to identify cell-type specific markers associated with patients’ survival along with the likely regulatory layer behind the differences in gene expression. We found that downregulation of proximal renal tubule genes and hence loss of cellular identity is implicated with stage IV patients’ survival. Here, we demonstrate how we can use SiRCle to uncover drivers across regulatory layers that may explain distinct cohorts of ccRCC, but we anticipate other cancers and data types can be analysed similarly.

## Results

### Functional effects of ccRCC are regulatory layer dependent

ccRCC is a heterogeneous cancer with transformation and progression linked to widespread dysregulation of biological processes. To understand how cellular processes are affected, analyses must extend across regulatory layers and therefore incorporate multiple omics’ datatypes. Recently, Clark *et. al*. and the CPTAC consortium, investigated the impact of genomic alterations on protein regulation and created revised subtypes based on their integrated analyses^14^. As part of their study, Clark *et. al*. generated protein, RNA, and DNA methylation measurements from 110 patients with ccRCC^14^. We sought extend their work by investigating the compounding effects of dysregulation by following gene changes from the DNA methylation via the mRNA to protein layers to unravel the regulatory layer of each gene. Since no DNA methylation samples from normal tissue were available, we coupled the CPTAC data with tumour adjacent normal (herein normal) DNA methylation data from 151 patients in TCGA^45^ (see **Supplementary Table 1** for sample annotations and Methods for quality controls). The CPTAC cohort is demographically homogenous, but includes cases across four stages of ccRCC and multiple mutational patterns (**Supplementary Table 1, Supplementary Fig. 1a)**. Data types are herein denoted “data layer” as they correspond to distinct layers of regulation (DNA methylation beta values, normalised mRNA expression and normalised protein expression). PCA established that the primary source of variation in a data layer (in isolation) is the sample type (tumour versus normal) with neither of the top two components explaining tumour stage, age, presence of a BAP1 or PBRM1 mutation (**Supplementary Fig. 1b)**.

We identified genes changing significantly between tumour and normal by performing differential analysis tests for each data layer and found widespread changes, most notably on the mRNA layer (Methods, **Supplementary Table 1**). While there is minimal negative correlation between gene changes on the DNA methylation and mRNA layers, the mRNA and protein layers are strongly correlated **(Fig. 1a)**. We found 689 significant genes shared across the mRNA and protein layers and 461 significant genes between all three layers (see Methods for thresholds).

**Fig. 1.**
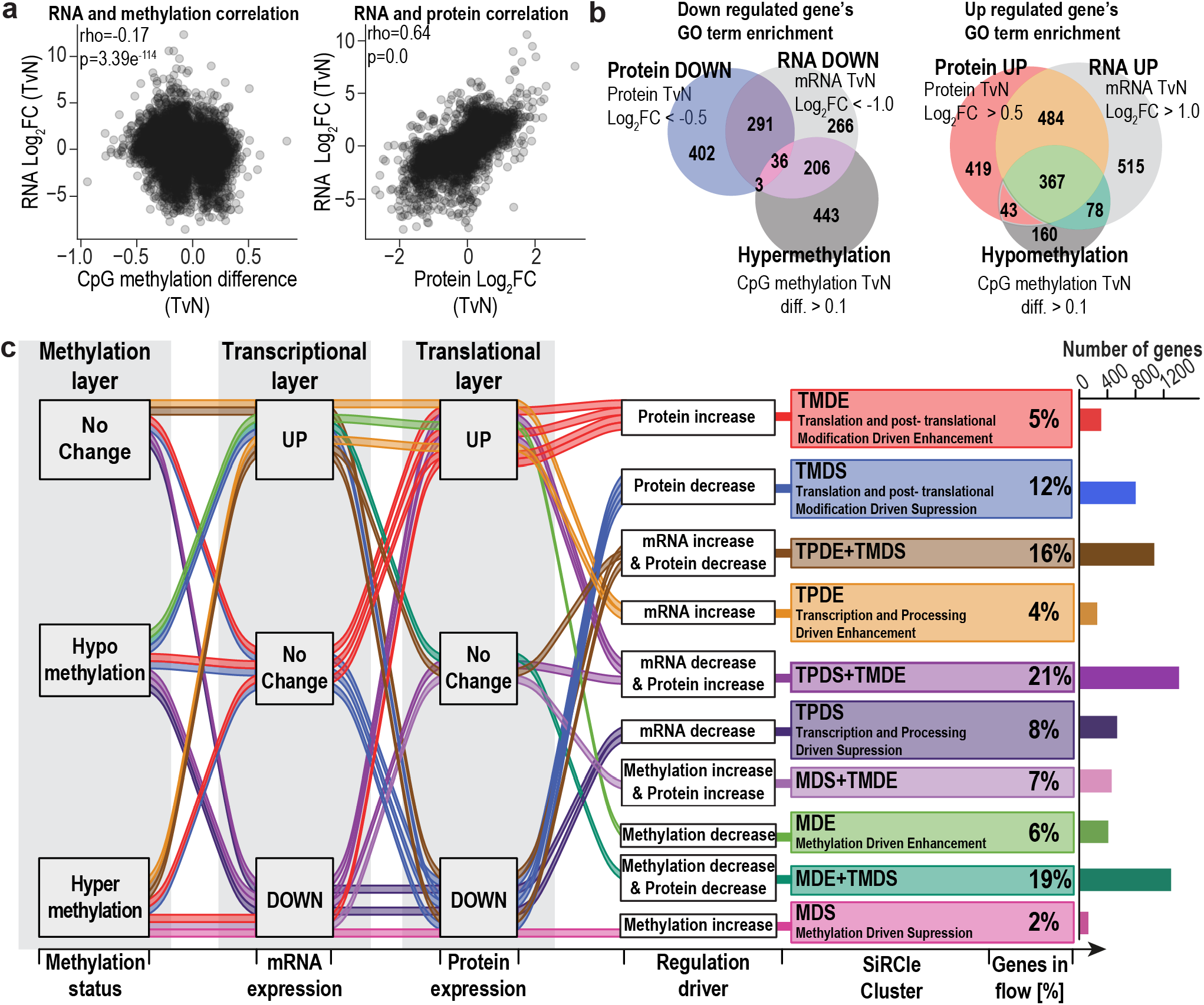
**a**, Spearman correlation comparing the change between tumour and normal samples for mRNA expression and DNA methylation, and mRNA expression and protein expression. **b**, Numbers of enriched Gene Ontology (GO) terms when comparing Tumour versus Normal (TvN) for each data type, indicating shared terms by overlapping regions in the Venn diagram. “Down regulated” corresponds to GO terms enriched in genes that are decreased in tumour samples, whilst “Up regulated” is for GO terms enriched in genes increased in tumour samples. **c**, Alluvial plot depicting the flows used to define SiRCle clusters. The plot is read from the left to the right, in line with the flow of information in a biological system. Each data type has been labelled as a layer, with one of three states defined for each layer based on the results for differential analysis between tumour versus normal in that data type. On the regulation driver we collapse the flows to include at most two regulation drivers, this we denote as “Grouping 2”, collapses are done on the later stage, for example, for TPDS+TMDE we have potential changes on all three layers, thus the first is collapsed. Direct flows correspond to relationships that follow the central dogma of biology, for example the flow followed in MDS. Overlaps in the Venn diagram are coloured according to the relationships in the alluvial plot that they capture, note not all relationships are presented in the Venn diagram.

Given the small number of significant genes shared across layers, yet comprehensive changes within each layer, we sought to determine whether the affected genes shared biological function. To do this, we performed Over Representation Analysis (ORA) of Gene Ontology (GO) terms on significant genes from each differential analysis, sub-setting genes by direction of change. We observed both unique and shared biological functions between the layers, with more similar functions enriched across layers for upregulated than downregulated genes **(Fig. 1b)**. Hypermethylated genes were most enriched for terms associated with development, while repressed genes were most associated with transporter activity (mRNA layer) and mitochondrial processes (protein layer) (**Supplementary Fig. 1c-d, Supplementary Table 2)**. In line with the findings from the original study^14^, immune response terms showed the most significant enrichment across all data layers for genes upregulated on the protein and mRNA layers (**Supplementary Fig. 1c-d, Supplementary Table 2)**.

### Following the flow of information extracts clusters that drive specific cellular phenotype in ccRCC patients

Given the lack of correlation between the data layers and the heterogeneity of functional enrichment resulting from them, we posited if grouping genes by their pattern across layers, prior to performing ORA would facilitate biological interpretation. Based on the differential analysis, we defined three states between tumour and normal, namely positive, negative or unchanged, for each gene and layer (**Fig. 1c, Table 1**, Methods section). To perform biologically meaningful clustering, we labelled each gene based on the regulatory layer where that gene is first “changed” in the tumour sample compared to the normal sample to bundle those following the same broad regulatory biology. We do this by following the central dogma of biology^46^, i.e. genetic information proceeds from DNA, to mRNA, and eventually protein (**Fig. 1c**). Since this is an ordered series of three 3-state transitions between the layers, there are 27 (3^3^) possible “flows”. For example, if a gene is hypermethylated, has a decrease in mRNA expression, and displays a decrease in protein expression, we can likely conclude dysregulation first occurred on the DNA methylation layer, adding this gene to the *Methylation-Driven Suppression* (MDS) cluster. While there are 27 possible flows, we define three granularities of clustering based on how many regulatory layers a user considers (Methods, **Table 1**). Throughout the paper, we use Grouping 2 with ten clusters defined by at maximum two changes, ensuring that for ccRCC genes are measured on the protein layer or on both the mRNA and DNA methylation layers **(Fig. 1c, Table 1)**. We term the process above “**Si**gnature **R**egulatory **Cl**ust**e**ring’’ (SiRCle).

**Table 1:** Regulatory labels from the different grouping methods.

| Methylation | RNA-seq | Proteomics | Regulation Grouping1 | Regulation Grouping2 | Regulation Grouping3 |
| --- | --- | --- | --- | --- | --- |
| Hypermethylation | DOWN | DOWN | MDS | MDS | MDS |
| Hypermethylation | UP | DOWN | TPDE+TMDS | TPDE+TMDS | TMDS |
| Hypermethylation | UP | UP | TPDE | TPDE | TPDE |
| Hypermethylation | DOWN | UP | MDS+TMDE | TMDE | TMDE |
| Hypermethylation | No Change | UP | TPDE+TMDE | TMDE | TMDE |
| Hypermethylation | No Change | DOWN | TPDE+TMDS | TMDS | TMDS |
| Hypermethylation | UP | No Change | TPDE+TMDS | TPDE+TMDS | TMDS |
| Hypermethylation | DOWN | No Change | MDS+TMDE | MDS+TMDE | TMDE |
| Hypermethylation | No Change | No Change | MDS-ncRNA | MDS_ncRNA | MDS_ncRNA |
| Hypomethylation | DOWN | DOWN | TPDS | TPDS | TPDS |
| Hypomethylation | UP | DOWN | MDE+TMDS | TMDS | TMDS |
| Hypomethylation | UP | UP | MDE | MDE | MDE |
| Hypomethylation | DOWN | UP | TPDS+TMDE | TPDS+TMDE | TMDE |
| Hypomethylation | No Change | UP | TPDS+TMDE | TMDE | TMDE |
| Hypomethylation | No Change | DOWN | TPDS+TMDS | TMDS | TMDS |
| Hypomethylation | UP | No Change | MDE+TMDS | MDE+TMDS | TMDS |
| Hypomethylation | DOWN | No Change | TPDS+TMDE | TPDS+TMDE | TMDE |
| Hypomethylation | No Change | No Change | MDS+ncRNA | MDE_ncRNA | MDE_ncRNA |
| No Change | DOWN | UP | TPDS+TMDE | TPDS+TMDE | TMDE |
| No Change | UP | DOWN | TPDE+TMDS | TPDE+TMDS | TMDS |
| No Change | DOWN | DOWN | TPDS | TPDS | TPDS |
| No Change | UP | UP | TPDE | TPDE | TPDE |
| No Change | No Change | UP | TMDE | TMDE | TMDE |
| No Change | No Change | DOWN | TMDS | TMDS | TMDS |
| No Change | UP | No Change | TPDE+TMDS | TPDE+TMDS | TMDS |
| No Change | DOWN | No Change | TPDS+TMDE | TPDS+TMDE | TMDE |
| No Change | No Change | No Change | No Change | No Change | No Change |

We performed ORA on the SiRCle clusters and found that each cluster was enriched for a clear biological signature important in ccRCC and consolidated the functional enrichment results we observed across each layer when independently analysed (**Fig. 2a, Supplementary Table 3**). Strikingly, we found clear biological signatures for both, SiRCle clusters regulated on one layer and SiRCle clusters regulated on multiple layers **(Fig. 1c)**. We found that hypomethylation (*Methylation-Driven enhancement* (MDE)) and enhanced transcription (*Transcription and Processing Driven Enhancement* (TPDE)) are associated with the hypoxic response, oxidative stress response and angiogenesis (**Fig. 2a**), key players in ccRCC tumour^42^. These terms were not revealed in the top terms when we performed ORA on each layer independently **(Supplementary Fig. 1c-d)**.

**Fig. 2.**
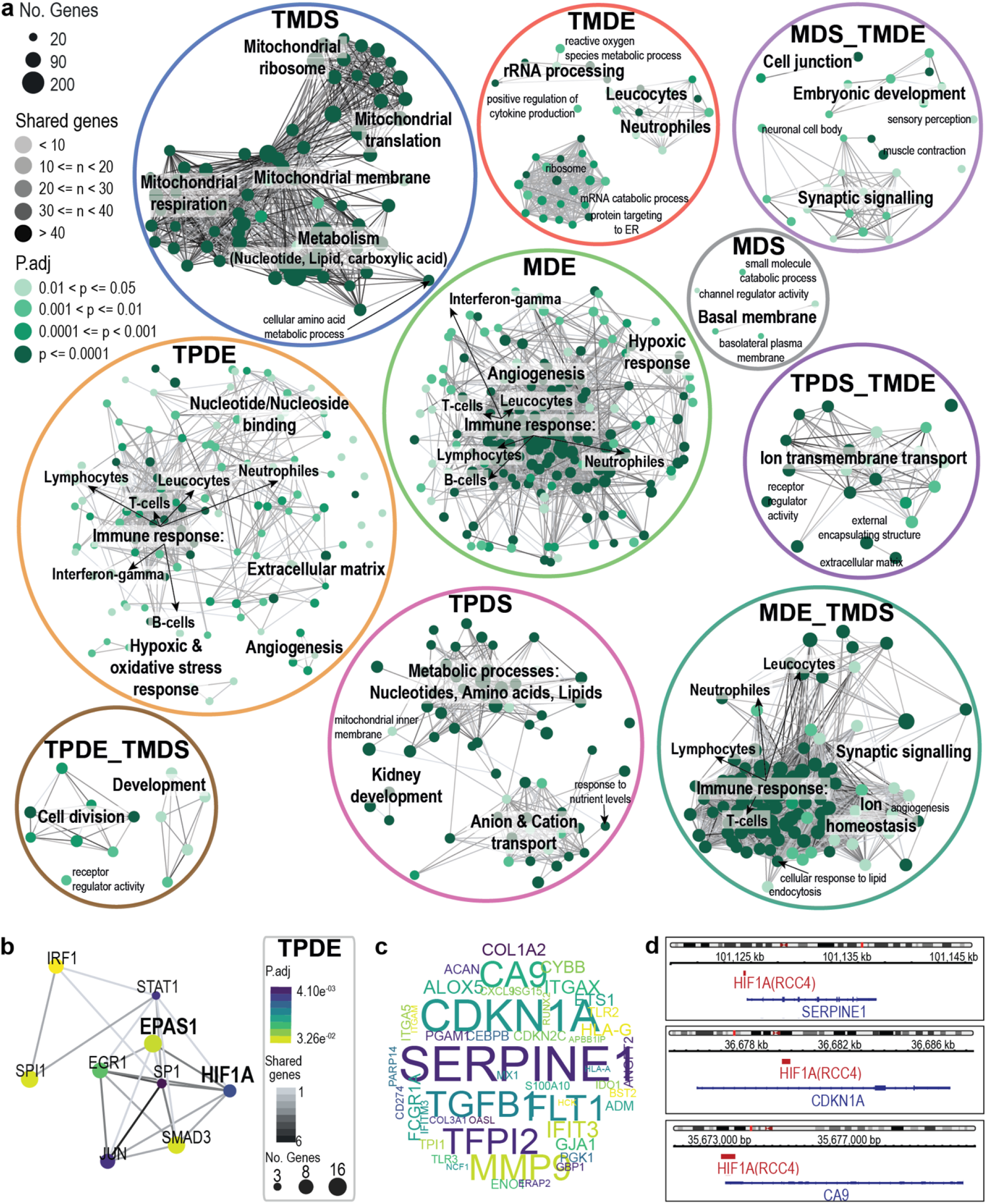
**a**, Emapplot visualisation of the over representation analysis (ORA) performed on each SiRCle cluster resulted in biological pathways that are altered comparing tumour versus normal. Pathways were plotted if p-adjusted value (p.adj) < 0.05 and the gene ratio was greater than 5%. The dot size corresponds to the number of genes found in the cluster that are part of the biological pathway. The colour of the dot shows the p.adj of the ORA. The connecting lines (grey) show that the biological pathways have genes in common. **b**, Transcription factor (TF) factor network of TFs from manually curated repositories (DoRothEA) that may drive genes in the cluster Transcription and Processing Driven Enhancement (TPDE). The dot size corresponds to the number of genes targeted by the TF. The colour of the dot shows the p.adj. The connecting lines (grey) shows the number of common genes the connected TFs regulate. **c**, Wordcloud including the TF targets with shared TF binding in the TPDE cluster, with the size corresponding to how many different TF’s are predicted to regulate the gene’s expression. **d**, HIF1A ChIP-seq peaks (RCC4, GSM3417827) binding at the transcription start site of *SERPINE1, CDKN1A* and *CA9*.

In line with previous findings^14,42,47^ we observed an increase in immune response pathways, and show these genes are likely regulated by DNA hypomethylation (MDE, **Fig. 2a**) or enhanced transcription (TPDE, **Fig. 2a**). Clark *et. al*. described a high correlation between mRNA and protein expression of immune signatures^14^, supporting our finding that the regulation occurs at the transcriptional layer. We were also able to distinguish between immune response genes likely regulated by hypomethylation from those likely regulated by enhanced transcription.

We observed that genes in two SiRCle clusters are involved in distinct metabolic rewiring, with metabolic processes such as lipid and amino acid metabolism likely regulated by transcriptional suppression (*Transcriptional and Processing-Driven Suppression* (TPDS)), whilst mitochondrial respiration and nucleotide metabolism are likely downregulated by translational suppression of the proteins (*Translation and translational-modification-Driven Suppression* (TMDS), **Fig. 2a**). Genes regulated by translational suppression appear to be involved in mitochondrial metabolism, mitochondrial translation and mitochondrial morphology (TMDS, **Fig. 2a**). Transcriptionally repressed genes were enriched for kidney development, hinting toward a loss of cellular identity in cancer cells (TPDS, **Fig. 2a**). Using SiRCle clustering we were able to determine the layer at which a gene’s dysregulation occurs and find clusters corresponding to distinct biological processes that may underly tumour pathology.

### Transcription factors as drivers of genes in SiRCle clusters

A subset of SiRCle clusters contain genes that change their state at the transcriptional (TPDE/TPDS) and/or the methylation layer (MDE/MDS), suggesting that they are regulated by transcription or epigenetic factors. Indeed, a specific TF may drive such changes in gene expression as TF’s can act to enhance/repress gene transcription and DNA methylation changes can alter the TF’s ability to bind to the DNA^48^. We used validated TF-to-target interactions from DoRothEA^49^ to identify what TFs were statistically associated with each SiRCle cluster. We use Fischer’s Exact Test (FET) to measure association between a TF and SiRCle clusters by testing if a TF targets more genes in a single cluster relative to background chance (all SiRCle clusters). This analysis only recovered TFs significantly associated with SiRCle clusters that are either up- or down-regulated at the methylation and transcriptional layers (**Fig. 2b, Supplementary Fig. 2a-b Supplementary Table 4)**. The stratification provided via SiRCle clustering helps distil the data from which regulatory logic can be inferred.

Given that HIF1 drives angiogenesis in ccRCC tumours^42^, we were unsurprised to find HIF1 TFs target genes in TPDE and MDE as both these clusters were enriched for angiogenesis related GO terms (**Fig. 2a-b, Supplementary Fig. 2a, Supplementary. Table 4**). We found that both HIF1A and EPAS1 (also known as *HIF1B*) had a significant increase in protein expression and were significantly associated with SiRCle clusters with an increase in mRNA and protein expression (TPDE, MDE). Given DoRothEA is not tissue specific we sought to confirm that the HIF TFs bind to the target genes in ccRCC by using ChIP-seq data from HIF1A and EPAS1 in kidney cancer cell lines (see Methods). We found evidence of HIF1A binding at the transcription start site (TSS) of genes in both TPDE (*SERPINE1, CDKN1A* and *CA9)*, and MDE (*VEGFA, EGFR* and *SLC2A1)* (**Fig. 2c, Supplementary Fig. 2d**). Similarly, we found evidence of EPAS1 binding at the TSS of *SERPINE1* and *CA9* in the TPDE cluster, confirming both HIFs ability to bind to target genes in ccRCC cell lines (**Supplementary Fig. 2e**). TF enrichment in SiRCle clusters elucidates changes to TF activity that affect transcriptional regulation at both the transcriptional and methylation layers.

### ccRCC characteristic metabolic changes are detected on distinct layers

Metabolic rewiring plays a crucial role in ccRCC and hence we sought to investigate the dysregulation of genes involved in metabolic rewiring in ccRCC to understand the layer at which the metabolic pathways are altered. In order to do so, we performed Gene Set Enrichment Analysis (GSEA) using metabolic signatures from Gaude *et. al*^50^, herein referred to as metabolic signatures, to identify pathways with coordinated protein changes in tumour versus normal. The protein layer was chosen for GSEA to capture changes in pathways’ enzyme activity, and then coupled with SiRCle cluster annotations to relate gene changes to the initial layer of dysregulation (**Supplementary Table 5**). We found the majority of enzymes involved in glycolysis (70%) were increased comparing tumours versus normal, fitting with the previously established upregulation of glycolysis in ccRCC^8^. Interestingly we find that some glycolytic enzymes are hypomethylated and upregulated on both the mRNA and protein layers (SiRCle cluster MDE, **Fig. 3a** and **Supplementary Fig. 3a**). From the TF analysis we noticed that H1F1A targets in MDE included several glycolytic enzymes, and we also found H1F1A binding at the transcription start site of glycolytic genes in kidney cancer cell line ChIP-seq data (**Supplementary Fig. 3c**).

**Fig. 3.**
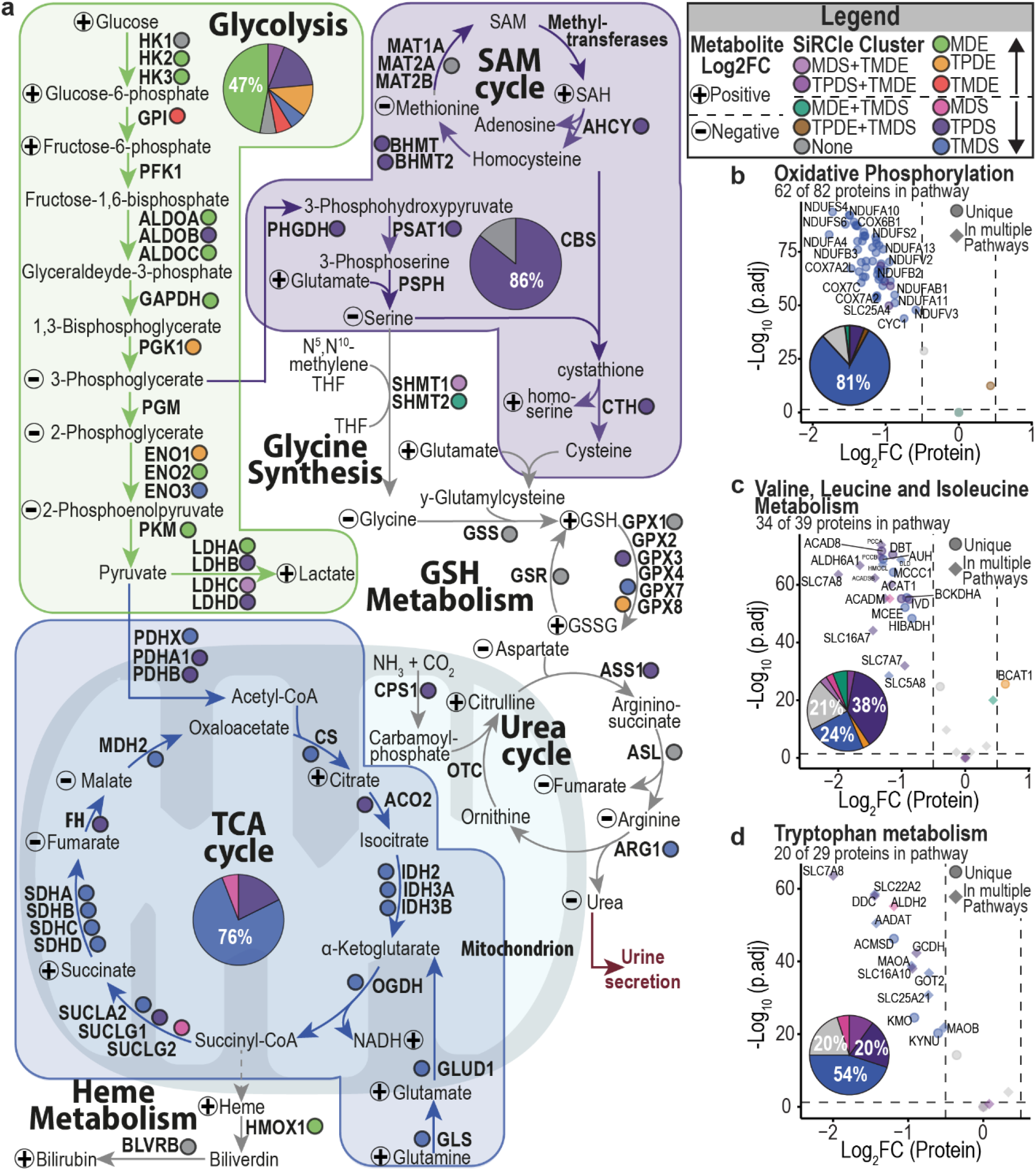
**a**, Metabolic changes of glycolysis, tricarbolic acid (TCA) cycle and serine, glycine and cysteine biosynthesis in ccRCC comparing tumour versus normal. Metabolic enzymes are written in bold and if they have been detected, they are labelled with coloured circles depending to the SiRCle cluster they are part of. Metabolic change of detected metabolites is labelled with plus for a positive and minus for a negative log_2_FC. For the three main metabolic sections are based on the SiRCle cluster the majority of metabolic enzymes are part of and are summarised in pie charts. **b-d**, Volcano plots are based on the metabolic pathways defined by Gaude *et. al*^50^ and the number of proteins detected of the pathway are reported. Proteins that are unique for the metabolic pathways are displayed in circles and proteins that are part of multiple metabolic pathways are displayed in diamond. The colour code is based on the SiRCle cluster the protein is part of and is summarised in the pie chart. GSEA was performed using the protein statistic. Here we plot oxidative phosphorylation **(b)**, valine, leucine and isoleucine metabolism **(c)** and tryptophan metabolism **(d)**, all reported a GSEA p.adj of 0.001518315.

We next sought to dissect the role of mitochondrial dysfunction in ccRCC, including the suppression of mitochondrial electron transport chain (ETC), since this has been previously confirmed^8^ to play a role in ccRCC, yet how metabolic enzymes are regulated remains unclear. We found that electron transport chain complexes were depleted in the tumour samples with the decreased expression of enzymes likely regulated at the translational layer (SiRCle cluster TMDS, **Fig. 3d**). The translation-driven downregulation affects almost all enzymes of the TCA cycle (SiRCle cluster TMDS, **Fig. 3a** and **Supplementary Fig. 3d**). In addition to these primary metabolic rewiring steps, there are many secondary pathways altered in ccRCC^8^. By coupling SiRCle annotations with GSEA we found that the suppression of cysteine methionine and glutathione (GSH) metabolism^7^ occur on the transcriptional layer (SiRCle cluster TPDS, **Fig. 3a** and **Supplementary Fig. 3e, f**). Similarly, serine biosynthesis, which is important to fuel GSH biosynthesis also occurs on the transcriptional layer. In accordance with the ORA results (**Fig. 2a**), we found that other amino acids’ metabolic pathways such as tryptophan, valine, leucine and isoleucine are downregulated either by translational or transcriptional suppression (SiRCle clusters TPDS and TMDS, **Fig.3d-f**).

To understand the impact of the enzyme expression on the metabolic profile, we used metabolomics data from 84 ccRCC patients with paired tumour and adjacent tissue samples^7^ and performed differential metabolomics analysis to identify the metabolites significantly changing between tumour and adjacent tissue. To consolidate the metabolite levels with enzyme information, we manually assigned significant metabolite changes to the pathways identified from the protein layer GSEA (**Fig. 3a**). We observed a “split” of glycolysis as previously described by Hakimi *et al*^7^, whereby metabolites upstream of GADPH are accumulated, and the downstream metabolites are depleted, despite the majority of glycolytic enzymes being upregulated on the protein layer (**Fig. 3a, Supplementary Fig. 3b-c, Supplementary Table 5**). A similar split is observed in the TCA cycle, where citrate and succinate are increased despite the decreased enzyme’s protein expression, whilst fumarate and malate are depleted in line with the decreased enzyme’s protein expression (**Fig. 3a**).

Overall, we show that SiRCle clustering can elucidate the layer where metabolic changes in ccRCC are orchestrated, highlighting points of intervention for targeting metabolism as anticancer strategy.

### Functional differences between late and early-stage tumours show limited agreement across layers

The previous sections explored dysregulation in tumour samples across a heterogeneous cohort of ccRCC patients. However, it has been reported that tumour stage is an important determinant of prognosis and treatment response^41^. We thus sought to identify the regulatory differences between patients with stage I and stage IV tumours by performing differential analysis on each layer, between tumour versus normal, for each patient group independently (**Supplementary Table 6**). Using the significant genes from each analysis, we performed ORA on each layer. On the protein and mRNA layers, we found that similar biological terms were enriched across the two patient’s groups, whilst on the methylation layer there were far more terms enriched in the stage IV samples. The main differences between the top associated GO terms were an association with extra cellular matrix terms in stage IV samples **(Supplementary Fig. 4a)**.

It is unsurprising that the tumour versus normal comparisons yielded similar results despite being run on patients with different stage tumours as the majority of variation in the dataset was accounted for by the sample type. Hence, we next tested for differences in tumour profiles between stage IV and stage I samples **(Supplementary Table 6)**. When comparing stage IV versus stage I tumours we found few changes on the protein layer, but aberrant changes on the DNA methylation and mRNA layers **(Supplementary Fig. 4b)**. As tumours progress, the dynamic adaptability of metabolism plays a crucial role to ensure cell growth^51^, and therefore, we tested for coordinated changes across metabolic signatures between using the differences between stage IV and stage I tumours. We found minimal shared enrichment of pathways between layers, with only N-Glycan Biosynthesis positively enriched on both the protein and mRNA layers, none shared on the mRNA and methylation layers, and Cyclic Nucleotides Metabolism, Glycolysis and Gluconeogenesis, Carbonic Acid Metabolism, shared between the negatively enriched protein layer and positively enriched methylation layer **(Supplementary Fig. 4c)**. While we find significantly changing metabolic pathways within each layer, functional information is not readily captured across layers rendering the results challenging to interpret.

### Novel integrated statistical test to identify changes between patient cohorts

Given the independently performed differential analyses were unable to extract shared functions when comparing patient’s with stage I versus stage IV tumours, we posited that integrating across the layers prior to performing differential analysis and biological enrichment may better capture the biological signal. We opted to use a variational autoencoder (VAE) to learn gene-wise relationships *across* the three data layers (**Supplementary Fig. 5a)**. Akin to matrix factorisation methods, a VAE finds a projection of data that centres on variance. However, unlike linear methods, such as PCA, a VAE does this using a neural network and can thus capture non-linear relationships.

In short, there are two parts of a VAE, 1) an encoding function that transforms data to a compressed representation, and 2) a decoding function that recreates the input from the compressed representation. The encoding and decoding functions are shared once the parameters have been learnt from the training data, while an encoded value is specific to a given datapoint. For example, the compressed representation of data point *x*_*ip*_ is given by: *z*_*ip*_ ≅ q_θ_(*z*∣*x*_*ip*_) ≅ *µ*_*ip*_ + *σ*_*ip*_ ⊙ *ϵ*_*l*_, where *z*_*ip*_ is the latent representation produced by the model conditioned on *x*_*ip*_, *q*_*θ*_ is the encoding function, *ϵ*_*l*_ is stochastic noise and part of regularisation, *µ*_*ip*_ is the encoded mean and *σ*_*ip*_ is the encoded variance. Note, *x*_*ip*_ refers to a feature vector of a gene, denoted by index *i* for a specific patient *p*. The feature vector is the normalised values across the data layers (**Supplementary Fig. 5a)**.

The encoding and decoding functions are “learnt” by minimising an objective lower bound over a dataset, by iteratively updating parameters over a number of epochs. For our purposes, the objective is to minimise the difference between input and output as per mean squared error and maximise the similarity between the latent space and a gaussian normal distribution via mean maximum discrepancy (see Methods for specifics).

Given the relatively low signal produced by stage, extracting regulatory variations may be obstructed by the noise in the dataset when considering all regulatory relationships at once. As such, instead of learning a dataset-wide representation, we used genes from each SiRCle cluster to learn a representation that defines a given regulatory flow. As such, there is an encoding function *q*_*θ*_(*z*|*x*) for each SiRCle cluster.

As we can now calculate an integrated (encoded) value for each gene, for each patient, we can define a gene’s *integrated difference* as a gene’s mean encoding difference between two sets of patients, e.g. patients with stage IV tumours versus stage I tumours (**Supplementary Fig. 5a**).

Mathematically we define this below, where *z* refers to the latent encoding given a particular data point, which corresponds to a patients’ gene value across the layers. Gene index *i* is held constant as the mean difference is calculated between patients sets, for example patients with a stage IV tumour may be in set *S*, and those with a stage I tumours are in set *G*.

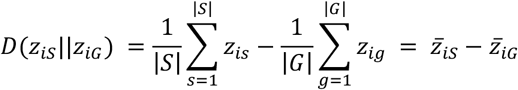

For biological interpretation, we can correlate the integrated difference to a biological reference such as the mean difference between the two patient sets on the protein layer. For example, in the MDE SiRCle cluster, we see that the stage IV versus stage I integrated difference correlates to the difference on the protein and mRNA layers between these patient sets (**Supplementary Fig. 5b, Supplementary Table 7**).

Using our integrated value for each gene, we next perform a Mann-Whitney U test to identify genes with a significant integrated difference between patients with early (stage I, N=45) and late (stage IV, N=10) stage tumours, old (N=56) and young (N=7) patients, and patients with a BAP1 (N=13) mutation or PBRM1 mutation (N=33) (**Supplementary Fig. 5b**). In the following sections, we demonstrate that VAE integration prior to performing analyses enables us to identify changes across the regulatory layers that were not found when performing differential analyses independently on each data type.

### Metabolic alterations in tumour stage and patients age impact on ccRCC’s metabolic fingerprint

While there were few shared metabolic pathways in the layer-specific analyses, an integrative model may improve our ability to understand metabolic differences between ccRCC patient’s groups. As metabolic profiles alter not only during tumour progression^51^, but also during ageing^52^, we performed two comparisons using the integrated values between 1) patients with stage I and stage IV tumours and 2) old and young patients. For each comparison, we performed GSEA using the metabolic signatures to identify pathways with coordinated integrated differences between two groups (**Fig. 3**). We define regulation as “up” if the integrated value increases, and as “down” if the integrated value decreases. Given the integrated value is positively correlated with the mRNA or protein layers, it carries biological meaning yet e.g. “up” regulation of the integrated value of a specific gene does not necessarily equal the same increase in mRNA or protein (**Supplementary Fig. 5b, Supplementary Table 7**). In line with previous observations^14^, oxidative phosphorylation was up regulated when comparing stage IV with stage I (**Fig. 4a, Supplementary Table 7**), yet we are now able to understand on what layer the enzyme expression is defined and observed that oxidative phosphorylation enzymes are likely altered on the translational layer. Reassuringly, oxidative phosphorylation was also a significant pathway in the stage IV versus stage I tumour protein layer GSEA analysis, but not on the mRNA layer, matching our TMDS label and showing that trivial single-layer relationships are captured by the VAE approach (**Supplementary Table 6**). Surprisingly, the coordinated change observed in oxidative phosphorylation was not observed for enzymes of the mitochondrial TCA cycle, with only some enzymes such as *IDH1, FH* and *PCK1* up in stage IV compared to stage I (**Fig. 4b**). Interestingly, citrate and cis-aconitate metabolite levels were depleted in late-stage compared to early-stage patients, whilst the downstream metabolites remained unchanged over the stages and have decreased protein expression in the tumour tissue (**Fig. 4c**). Oxidative phosphorylation showed a more variable, albeit similar trend in old patients when compared to young patients (**Supplementary Fig. 6a, Supplementary Table 7)**. Conversely, TCA cycle enzymes were significantly up in old patients compared to young patients when performing GSEA and accordingly, metabolite levels were increased for succinate, fumarate, and malate in old patients (**Supplementary Fig. 6b, Fig. 6c**). Noteworthy, the majority of old patients do not present late-stage tumours (old patients: stage I = 27 (41%), stage II = 9 (14%), stage III = 24 (36%), stage IV= 6 (9%); young patients: stage I = 7 (78%), stage 2 = 1 (11%), stage 3 = 1 (11%) **Supplementary Table 1**).

**Fig. 4.**
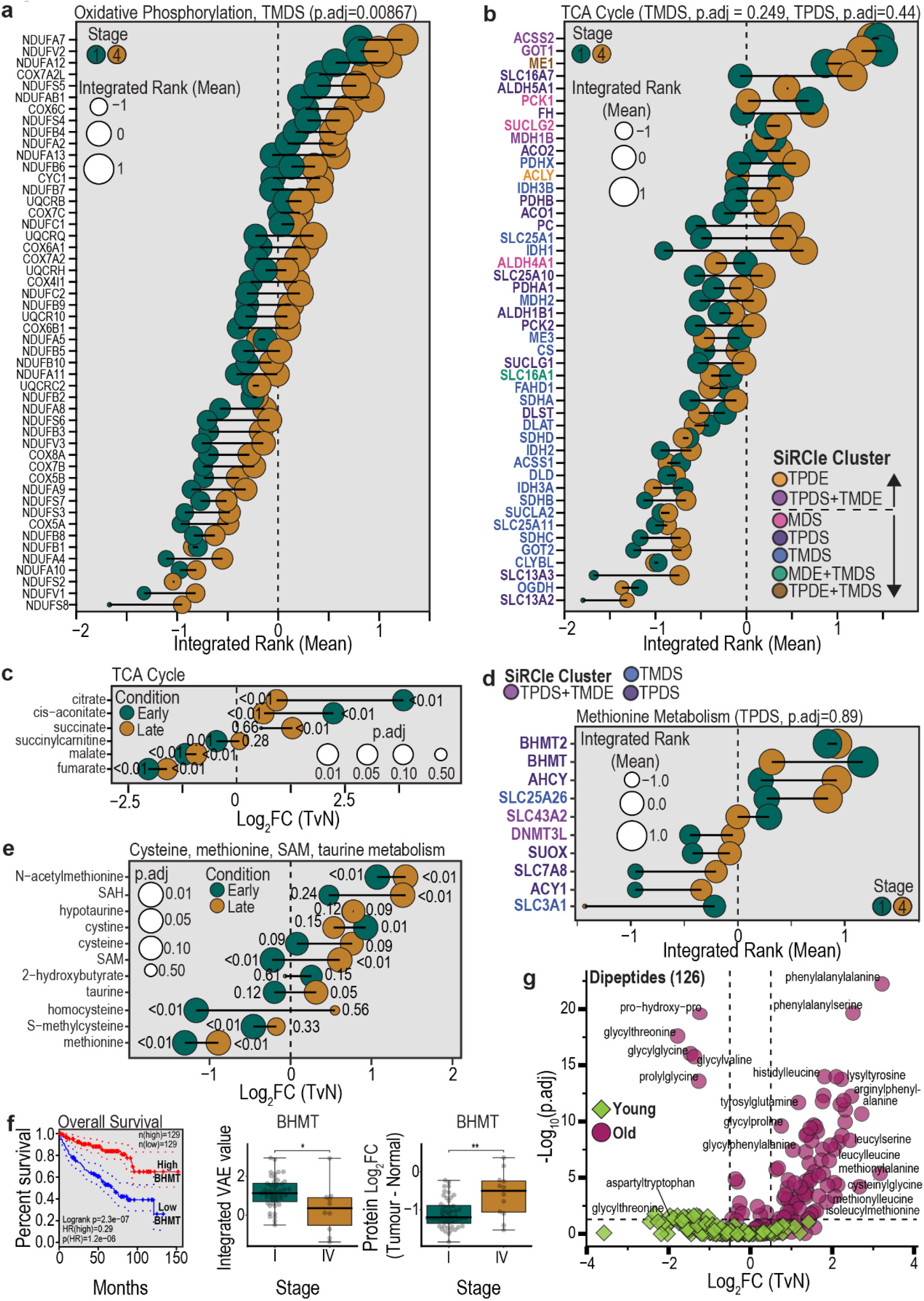
Metabolic pathways based on gene expression are defined by Gaude *et. al*^50^ with p.adj values corresponding to the GSEA results after ranking the genes in each SiRCle cluster using the VAE integrated rank. Metabolite pathways based on metabolites are defined by Hakimi *et. al*^7^. **a**, Comparison of the VAE integrated rank of stage IV and stage I patients for oxidative phosphorylation for genes within the Translation and post-transcriptional Modification Driven Suppression (TMDS) SiRCle cluster. **b**, Comparison of the VAE integrated rank of stage IV with stage I patients for TCA cycle genes colour coded for the SiRCle clusters the gene belongs to. **c**, Comparison of the Log2FC of tumour versus normal (TvN) of late stage (III & IV) with early stage (I & II) patients for TCA cycle metabolites. **d**, Comparison of the VAE integrated value of stage IV with stage I patients for genes corresponding to the methionine metabolism pathway colour coded for the SiRCle clusters the gene belongs to. **e**, Comparison of the Log2FC (TvN) of late stage (III & IV) with early stage (I & II) patients for “cysteine, methionine SAM, taurine metabolism” metabolites. **f**, BHMT survival curve based on (http://gepia2.cancer-pku.cn/) using quantile default cut-offs of 25%, 0.75%. VAE integrated rank alongside protein Log2FC (TvN) for stage I with stage IV patients. **g**, Comparison of the Log2FC (TvN) of late stage (III & IV) with early stage (I & II) patients for metabolites of the dipeptide group.

A novel finding from Hakimi *et. al*. was that metabolites involved in methionine metabolism are altered between different tumour stages in ccRCC^7^. We sought to build on this by investigating whether the changes in methionine metabolism could be traced back to a regulatory change. We found that enzymes of the methionine metabolism are predominantly up on the transcriptional layer when comparing stage IV vs stage I (**Fig. 4d-e, Supplementary Table 3, Supplementary Fig. 4b)**. *BHMT, SLC43A2* and *SLC3A1*, were down in stage IV patients, while all other enzymes were up (**Fig. 4d**). We observed an overall accumulation of metabolites within this pathway in the late-stage patients, indicating methionine depletion is less severe in late-stage patients (**Fig. 4e**). It was surprising that despite the fact that BHMT is down in stage IV patients, metabolite levels are unaffected; this hinted towards a different role of BHMT in late-stage tumours (**Fig. 4d-e**). Given BHMT is a proximal tubule-specific protein and SLC3A1 is a kidney amino acid transporter mainly found in proximal tubule of the kidney, a loss of expression of BHMT and SLC3A1 may promote a loss of cellular identity in stage IV tumours, which in turn is detrimental for the patient’s survival (**Fig. 4f and Supplementary Fig. 6f**). Furthermore, it has been recently discussed that enzymes involved in methionine metabolism play a role in ccRCC patients’ survival^53^. We also found the methionine pathway to be affected when comparing old with young patients. In old patients all enzymes were consistently up, which is in line with the increased metabolite levels (**Supplementary Fig. 6d-e**). Methionine upregulation could increase the methylation potential and hence alter the DNA methylation landscape within a tumour. Finally, the most altered metabolic pathway from the differential metabolite analysis when comparing old versus young patients were dipeptides (**Fig. 4g**). Dipeptides are a nutrient signalling mechanism that is required for chronic myeloid leukaemia stem cell activity in vivo^54^ and have recently been proposed as novel cancer biomarkers^55^. However, given dipeptides are produced from polypeptides by dipeptidyl peptidase enzymes, which is a class containing many enzymes, it is difficult to connect specific enzymes, and thus layers of regulation, to this observation.

Together these results showcased the importance of taking patients’ information into account when analysing and interpreting the data and that VAE integration enables the comparison of small patient groups whilst considering differences across the methylation, mRNA and protein layers.

### Genes distinguishing patient subpopulations associate with distinct immune cell types and define patient’s survival

Investigating regulatory changes across a tissue only provides an overview of changes occurring within the numerous cell types that make up the tissue. However, given the challenges associated with epigenetic profiling at the single-cell level, it remains a cost-effective, and reliable approach. Moreover, broad changes in regulatory patterns may indicate a change in the composition of cell types within a tissue. Given that immune response plays a critical role in ccRCC and is associated with specific mutational patterns^42^, we sought to assess whether perturbations across regulatory layers between patient subpopulations are associated with specific cell types. We used scRNA-seq data from eight patients with advanced renal cell carcinoma from the multiregional study of Krishna *et al*., which identified 31 cell types^47^. Again, we divided SiRCle clusters (MDE, TPDE, TMDE, MDS, TPDS and TMDS) into up and down subsets of genes based on the gene’s VAE integrated difference between two patient groups, and then tested if these genes are overrepresented in a specific cell type.

Focusing on comparing stage IV versus stage I, we found that the broad cell types identified by Clark et al^14^, are recovered, namely, CD8A+ Proliferating and Exhausted (CD8+ inflamed), CD45-PAX8+ renal epithelium (CD8-Inflamed), CD45-Vascular Endothelium (VEGFA) (**Fig. 5a**). Moreover, it becomes clear that some SiRCle clusters are associated with specific cell types (**Fig. 5a**).

**Fig. 5.**
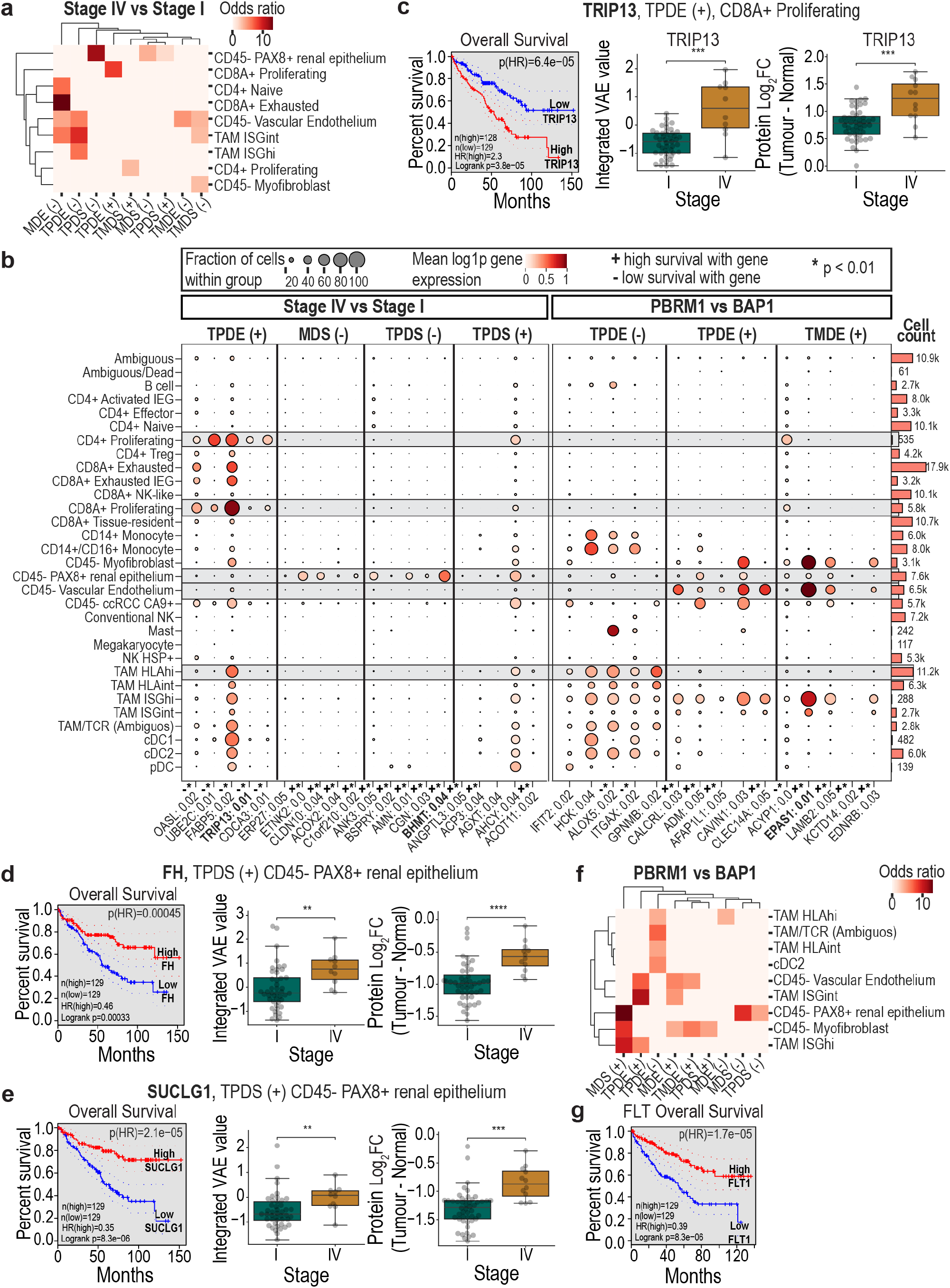
Boxplots show the integrated VAE rank for each patient in a subpopulation, alongside the protein log_2_FC between tumour versus normal (TvN) for that same group of patients. Survival plots were plotted using GEPIA2 (http://gepia2.cancer-pku.cn/). Differentially Expressed (DE) in the single cell clusters are genes that were significant (p < 0.05) in a one versus rest comparison and had a log_2_FC > 1.0 for a given cell type.**a**, Significant enrichment (odds ratio), using the Fischer’s Exact test of SiRCle clusters subset into (+) for significant (p <0.05) changes with stage IV > stage I, and (-) stage IV < stage I. **b**, Top 5 genes by expression in a specific cell type for SiRCle subsets along with summarised survival information from the Protein Atlas (https://www.proteinatlas.org/). Grey blocks highlight cell types discussed in the results. Similarly, with bold gene names. **c**, Survival and boxplots for TRIP13, which is in the Transcription and Processing Driven Enhancement (TPDE) (+) SiRCle cluster and CD8+ proliferating. **d**, Survival and boxplots for *FH*, which is in the Transcription and Processing Driven Suppression (TPDS) (+) SiRCle cluster and DE in CD45-PAX8+ renal epithelium. **e**, Survival and boxplots for *SUCLG1* which is in the TPDS (+) SiRCle cluster and DE in CD45-PAX8+ renal epithelium. **f**, Significant enrichment (odds ratio), using the Fischer’s Exact test of SiRCle clusters subset into (+) for significant (p <0.05) VAE integrated changes with PBRM1 > BAP1, and (-) PBRM1 < BAP1. **g**, Survival plot for *FLT1* from PBRM1 vs BAP1 comparison.

We found genes up in stage IV patients and regulated on the transcriptional layer (TPDE) that are enriched in CD8A+ proliferating cells (**Fig. 5a-b and Supplementary Fig. 6a**). This is in line with previous studies, which observed an overall increase in CD8+ T-cell in tumour-infiltrated immune cells in ccRCC compared to normal renal tissue^56^. Specifically, we saw enrichment of *TRIP13*, which has previously been associated with CD8+ T-cells^57^ and genome instability^58^. Reassuringly, we detected *TRIP13* as part of the TPDE up cluster and CD8+ enriched cell type and found that its expression is highly altered between stage I and stage IV patients and implicated in patient’s survival (**Fig. 5c**).

The CD45 PAX8+ renal epithelium cell type is specific for ccRCC^59^, hence we posited that genes within this group would be changed as the tumour progressed and lost its renal identity. Genes in CD45 PAX8+ renal epithelium were mainly further repressed in stage IV patients on the methylation and transcriptional layers (MDS down, TPDS down); however, we also saw recovery of expression in stage IV tumours on the transcriptional layer (TPDS up) (**Fig 5a-b and Supplementary Fig. 6b-d**). While extracellular terms were also observed in the differential analysis, these were only identified as top terms on the mRNA layer; given we observed the genes in TPDS, the changes are likely to translate to the protein layer **(Supplementary Fig. 4c)**. Metabolic enzyme BHMT is further repressed in late-stage patients (TPDS) further confirming its importance in the renal epithelium and that loss of expression may be associated with loss of cellular identity resulting in poor survival (**Fig. 4f**). Moreover, several mitochondrial enzymes up in stage IV patients (TPDS) are also associated with the renal epithelium, supporting evidence of complex metabolic rewiring in late stage renal tumours. Indeed, we found both *FH* and *SUCLG1* are associated with survival, indicating in late stage tumours, there is some recovery of these important enzymes (**Fig. 5d-e**).

Previous work has shown that the immune response plays a critical role in ccRCC and that germline mutations can affect ccRCC patients’ outcomes^12^. In addition to *VHL* mutations, *PBRM1* and *BAP1* facilitate the development of ccRCC^12^, however with differing outcomes where *BAP1* mutations are associated with a poor prognosis whilst *PBRM1* mutations result in less aggressive cancers^42^. Thus, we wanted to test if the differences in the regulatory profiles between *PBRM1* versus *BAP1* patients could explain the different patient outcomes. Hence, we used our VAE integrative comparison approach to identify genes with regulatory perturbations between patients with a PBRM1 mutation compared with patients with a *BAP1* mutation. We found genes changing significantly between the patients across all regulatory layers, and term the comparison *PBRM1* versus *BAP1* **(Supplementary Table 7)**. Interestingly, we found similar cell types enriched in TPDE up when comparing *PBRM1* with *BAP1* patients as in the TPDE down group from stage IV vs stage I patients (**Fig. 5a**,**f**), fitting with the association between *BAP1* having a generally poorer outcome^42^. TPDE and MDE down enrich for Tumour Associated Macrophages (TAMs) cell types, with the classically immunosuppressive cell cluster (TAM HLAhi), showing higher expression in *BAP1* (TPDE down) in both SiRCle clusters (**Fig. 5b, 5f**). In line with Clark *et. al*.*’s* finding that PBRM1 mutations associate with an angiogenesis immune population^14^, we observed an enrichment of genes up in *PBRM1* in CD45-Vascular Endothelium, which is connected to genes up in the SiRCle clusters MDE, TPDE, and TMDE (**Fig. 5f** and **Supplementary Fig. 6e**). Indeed, the latter included *EPAS1*, also known as *HIF2A*, and is an established prognostic marker in renal cancer with low expression correlating with poor prognosis^60^ (**Fig. 5b, Supplementary Fig. 6e**). Another known gene important in angiogenesis, *FLT1*, is also connected to the CD45-Vascular Endothelium and higher in *PRBM1* mutations (**Fig. 5b, Supplementary Fig. 6f**). *FLT1* is up in TPDE and low expression of *FLT1* is associated with poor prognosis (**Fig. 5g**). Interestingly, in our TF analysis earlier, we identified that *FLT1* was targeted by EPAS1, meaning that the increased expression of EPAS1 could be regulating the increased expression of *FLT1*; however, this would need to be confirmed in ccRCC patients as in cell lines we did not observe EPAS1 binding sites at *FLT1*.

Our results suggest that while single-cell multi-omic data is not readily available, or remains too costly, we can retrospectively look for the enrichment of genes with shared regulatory behaviour in scRNA-seq data to predict the likely cell types that will be affected in a given population and the layer of dysregulation behind the gene’s changes. This additional information provided by the SiRCle analysis can thus inform future multi-omic experimental decisions at the single cell level to further understand disease pathology.

## Discussion

Here we present SiRCle, a method for integrating DNA methylation, RNA-seq and proteomics data to extract support for what layer or layers (DNA methylation, transcription and/or translation) genes are regulated and/or perturbed between patient subsets with different phenotypic traits (**Fig. 6**). While SiRCle extends to any cancer, we applied it to clear cell renal cell carcinoma (ccRCC) patients. ccRCC is a particularly heterogeneous cancer with cascades of mutations and complex remodelling of the tumour microenvironment that affect several regulatory networks^3^, hence we expect many regulatory relationships to hinge on joint consideration of multiple data modalities.

**Fig. 6.**
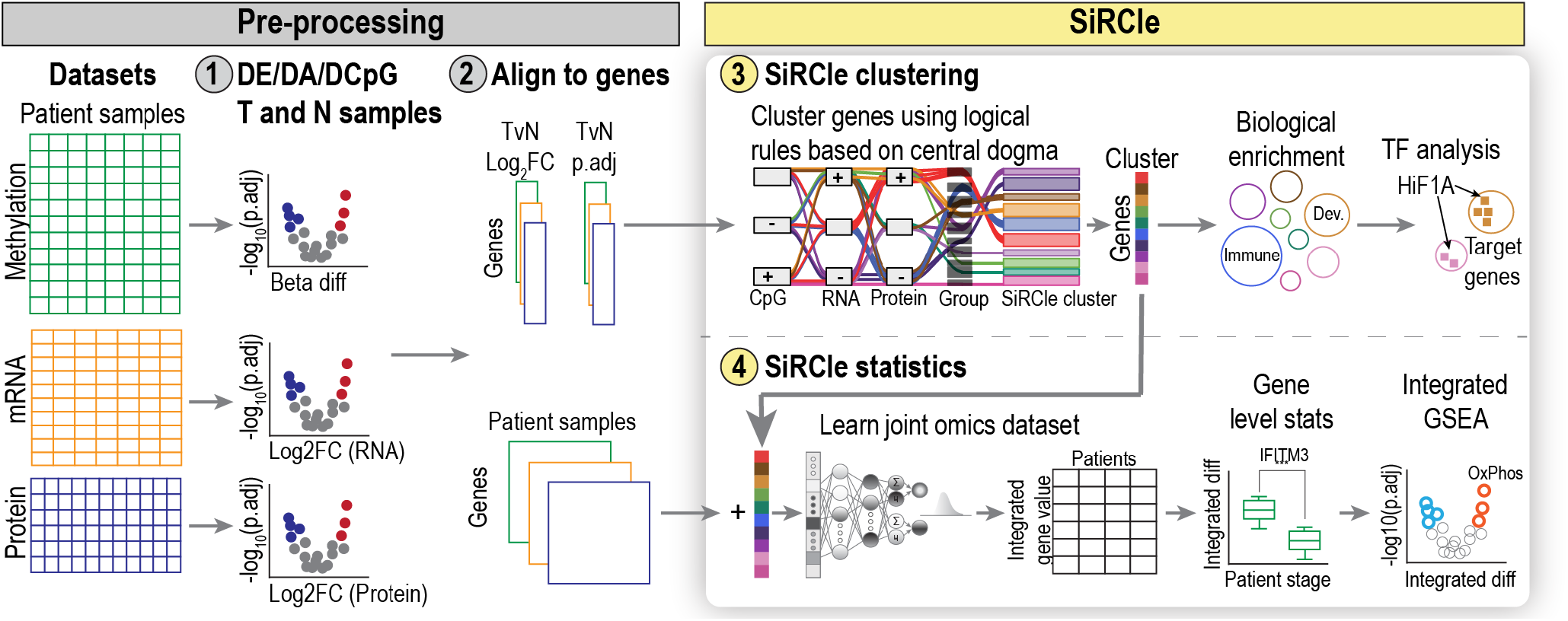
Overview of SiRCle (**<u>Si</u>**gnature **<u>R</u>e**gulatory **<u>Cl</u>**ust**<u>e</u>**ring).

Across a cohort of ccRCC patients, we show modest correlation between data at different layers as well as limited agreement among biological terms extracted from each layer in isolation. This observation underscores the challenges of analyses applied to individual omics since some of the gene changes observed at the epigenetic or transcriptional layer may not be translated into protein changes. We thus grouped genes based on regulatory “flows” based on differences (between control and disease) at each layer; we found the majority of genes exhibit states counter to the generalisation that increased DNA methylation at the gene represses transcription, which in turn decreases translation of the gene product within the condition, *or* (conversely) that reduced methylation at the gene activates transcription, which in turn increases translation of its product. While the extent of dissonance highlights the importance of considering multiple layers of regulation, we found that by grouping them by cross-layer “flows” instead elicited distinct biological terms, e.g. metabolism and immune response, subsuming the function extracted from each layer in isolation. The advantage of SiRCle clustering is that by considering all possibilities of regulation, we can pinpoint the likely layer at which genes are dysregulated in disease.

In addition to consolidating biological function across the layers, SiRCle clustering captured relationships from adjunct data, such as TF relationships and metabolic changes. We found TF’s targets fell predominantly in clusters regulated on the layer of transcription or methylation and were able to distinguish HIF TF targets that increased via transcription, from those that increased via hypomethylation. The advantage of performing the regulatory clustering prior to the TF analysis is that we can exclude TFs that are predicted to drive genes in clusters regulated at the layer of translation (e.g. TMDE, TMDS), given TFs work at the layer of transcription.

Coupling SiRCle annotations with metabolic activity, we were able to establish a more detailed view of how changes in regulation correspond to changes in metabolic pathways. We found that glycolysis was regulated on the methylation layer (MDE), whilst the TCA cycle and oxidative phosphorylation were regulated translationally (TMDS). We noticed that glycolytic enzymes are upregulated in the context of ccRCC, suggesting hypomethylation increases DNA accessibility^48^, potentially improving the capacity for H1F1A to bind and upregulate glycolytic enzyme transcription and eventual translation. Hakimi *et. al*. showed that metabolites of the glycolysis pathway split at the level of GAPDH, whereby metabolites upstream of this enzyme are accumulated, and the ones downstream are depleted^7^. We found that this is not in line with the enzyme expression, yet this counterintuitive finding could be explained by the fact that the catalytic function of many enzymes is altered by post-translational modifications such as phosphorylation (e.g. PGK1^61^ and PKM^62^) or oxidation (e.g. GAPDH^63^). In fact, it has been shown for the CPTAC patient cohort that *PGK1* and *PKM* have increased phosphorylation when comparing tumour versus normal^14^. Moreover, glycolytic flux towards lactate in ccRCC is inhibited by FBP1^64^, yet FBP1 is not mutated in this patient cohort.

Until this point, we investigated changes across the whole ccRCC cohort, however, there are distinct subpopulations of patients, with differing prognoses, for example, those with stage I or stage IV tumours. We sought to investigate the regulatory differences between patient subpopulations, however, when we analysed each layer independently there was limited overlap between the mRNA and protein layers. Thus, to find regulatory differences between patient subpopulations (e.g. age, stage, mutation), we developed a novel integration approach using a VAE to learn a single key feature from the DNA methylation, RNA, and protein layers for each SiRCle cluster. By integrating across the layers prior to performing statistical analyses we found significant differences in genes important for survival when comparing early and late stage patients and showed that the VAE results in biologically meaningful integrated values. Moreover, we showed that our integrative approach can be used to perform GSEA between patients’ groups.

SiRCle allowed us to identify key differences between patients’ subpopulations in terms of metabolic rewiring and the different cell types. Interestingly, oxidative phosphorylation was less suppressed in late stage and older patients, despite old patients being distributed across multiple cancer stages. We also observed that within the TCA cycle intermediates, citrate and aconitate were depleted in late-stage tumours, despite the increased expression of the enzymes involved in their biosynthesis. Whether this result indicates a further functional regulation of the TCA at the post-translational layer, as for instance PDH phosphorylation, which in turn controls citrate biosynthesis, needs to be further investigated. In addition, we observed an increase in succinate in late-stage patients and this finding could indicate that these tumours may be more hypoxic, considering the role of succinate as marker of low oxygen in tissue^65^. The increase in succinate could also be due to the need to generate NADH, where the TCA cycle fuels into succinate and the heme biosynthesis pathway as previously described^66^. In line with this, HMOX1 is upregulated, likely via hypomethylation, and the metabolites heme and bilirubin are also accumulated (**Fig. 3a**). As discussed by Hakimi *et al*, the TCA cycle could be fuelled via glutamate which fuels into citrate via reductive carboxylation^7^. Lastly, in line with the enzyme expression, serine levels are depleted (**Fig. 3a**). While similarities in metabolic profiles between the old and late-stage patients were identified, we also found differences in methionine metabolism. Indeed, we found that BHMT and SLC3A1 are further repressed in late-stage patients yet show increased expression in old patients. Importantly, high expression of these genes is crucial for patient survival. We also found BHMT was specifically expressed in CD45-PAX8+ renal epithelium cells pointing to a complex interplay between metabolic rewiring and cellular identity that indicates BHMT could be used as marker of cellular differentiation.

Checking the occurrence of specific mutations, which can have an impact of patient outcome^12^, we uncover that genes highly expressed in patients with BAP1 mutations are selectively expressed in tumour-associated macrophages, and indeed are negatively associated with patient survival. Interestingly, we found that the TF EPAS1, also known as HIF2A, could regulate the expression of genes enhanced in PBRM1-mutated tumours. However, further studies are needed to investigate the link between BAP1 and EPAS1/HIF2A. Given the expense and technical challenges with single cell multi-omics data, we expect the capability to retrospectively examine SiRCle cluster activity in scRNA-seq data to be of value for understanding gene dysregulation in the context of heterogenous cancers.

While we have focussed on ccRCC, with the increasing efforts to measure the proteome of cancer cohorts and other diseases the SiRCle method will be broadly applicable to investigating other datasets. Using SiRCle researchers can extend known relationships, find regulators and drivers of gene clusters, and extract genes that differ between patient cohorts in many different cancer types.

## Methods

### RNA-seq

Downloaded the data from TCGA hg38 gene expression and clinical patient data from the CPTAC kidney study. Six samples were removed due to low correlation (ρ < 0.75) were removed (**Supplementary Table 1**)

### Clinical

Mutation data were downloaded for the patients from the RNA-seq dataset using TCGA’s GDC client (https://gdc.cancer.gov/access-data/gdc-data-transfer-tool, version 1.6.1) and scidat (https://github.com/ArianeMora/scidat,v version 1.0.3). The mutation data were merged with the clinical information from CPTAC patients along with the CIMP-status_Chr-stability_supp-download file also from the CPTAC study. Patient age was grouped into three categories (<42 = young, 42 = middle, and >58 = old), based on information pertaining to pre-post menopause, instance rate and age of onset from sporadic as compared with inherited mutations^35–37,67^.

### Proteomics

Data were downloaded from the CPTAC data portal on 14^th^ of April 2021 (https://pdc.cancer.gov/study-summary/S050). 11,355 proteins were measured for 201 samples. Missing protein data were imputed using DreamAI ensemble method^68^ (https://github.com/WangLab-MSSM/DreamAI) Samples exhibited a high correlation (∼96%) post imputation, with tumour and normal samples clustering distinctly, and as such no protein samples were removed. The supplied protein names were mapped to hgnc symbols using biomart mappings (scibiomart, 1.0.2, https://github.com/ArianeMora/scibiomart), and for those without direct mappings (211 protein names) were mapped using the external_synonym. Afterwards, 11 protein names remained unmapped, namely APOBEC3A_B, ELOA3D, FLJ44635, GAGE2D, GPR75-ASB3, KIAA0754, KIAA1107, LOC110384692, PALM2, WRB-SH3BGR and ZNF664-RFLNA.

### DNA Methylation

Tumour DNA methylation files were downloaded from the CPTAC portal, under metadata (https://cptac-data-portal.georgetown.edu/study-summary/S050), methylation samples annotated as not ccRCC by the CPTAC authors were omitted from our analyses (C3L_00011, C3L_00010, C3L_00097, C3L_00096, C3L_00004, C3L_00079), leaving 104 ccRCC tumour methylation samples. These were merged by reading in each file before merging on CpG ids and retaining the Beta values for each CpG. Correlation between samples was computed with three tumour samples identified as outliers with low correlation ρ < 0.75 namely C3L-01882, C3L-01885, and C3L-01281, which were removed. Using the provided GencodeV12 annotation (Hg37) unannotated Loci were removed leaving 586,055 Loci. To ensure compatibility with the Protein and RNA-seq data that use Hg38 gene identifiers, we mapped the transcript IDs (Hg37) obtained from the DNA methylation data to ensembl gene IDs shared in both versions (Hg37 & Hg38). For transcripts without a direct mapping, we sought to map on the “external_gene_name”, “hgnc_symbol”, and “external_synonym” from the column “GencodeCompV12_Name” to identify the Hg38 ensembl ID. Only 38,352 CpGs remained unassigned, corresponding to 7,910 unique genes “GencodeBasicV12_Name”, which were removed. The remaining 547,703 CpGs were used for subsequent analyses and annotated to 27037 unique Hg38 ensembl IDs. Normal DNA methylation data were downloaded from the TCGA data portal, by selecting TCGA-KIRC patients, and solid tissue normal m450 DNA methylation beta values (level 3 files), the subsequent manifest file and clinical data were provided as input to scidat. Using scidat the CpG beta value files and patient mutation data were downloaded. TCGA CpG data were merged on the CpG identifier (composite element reference) with the gene annotated CPTAC DNA methylation data, giving 268,355 CpGs across the two datasets with 151 normal samples from TCGA and 100 tumour samples from CPTAC.

### Dataset generation

Dataset summaries: RNA-seq (66 normal and 173 tumour samples), Protein data (81 normal & 103 tumour samples), and CpG samples (151 normal & 100 tumour samples). PCA of samples in each dataset shows a clear distinction between normal and tumour samples **(Supplementary Fig. 1b)**.

### Differential analysis

Differential expression analysis on RNA-seq data was performed using DEseq2^69^ (version 1.32.0), using condition ID (tumour) as the factor. Genes were excluded if there were less than 10 samples with at least 10 counts in a gene resulting in 25,655 genes for DE analysis. Differential abundance analysis on proteomics data was performed using limma^70^ (version 3.48.3). Only the condition (i.e. Tumour or Normal) was used in the design matrix. lmfit with eBayes was used and the p values were corrected using the FDR method. For the methylation data (268,355), CpGs with low (mean CpG value < 0.05) or high mean methylation (> 0.95) were removed prior to performing the differential test, resulting in 195,195, and 187,770 CpGs. CpG M values, 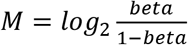 ^71^, were used as input to limma’s lmFit eBayes and p values were adjusted using the FDR method.

### SiRCle (<u>Si</u>gnature <u>R</u>egulatory <u>Cl</u>ust<u>e</u>ring)

SiRCle requires a single DNA methylation value to be assigned to a gene. After performing the CpG analysis using limma, CpGs were filtered in order to map only one CpG to a given gene. This was done by grouping the CpG data by annotated genes. In detail, if there were less than three significant (*p* < 0.05) CpGs, then the CpG with the greatest absolute change was selected. If there were more than 3 CpGs (with *p* < 0.05), we tested if the majority (> 60%) of CpGs agreed in the “direction of change” (positive or negative) and if not all CpGs were omitted. Otherwise, the CpG with the highest methylation difference was chosen.

Next, we performed the SiRCle clustering on the filtered CpG data, DE results from DEseq2 (RNA-seq Data) and the output from limma’s abundance analysis (Proteomics Data). The datasets were merged using the background method: (P&M)|(P&R)|(M&R) meaning a gene was retained if it was changed significantly (*p* < 0.05) in at least two of the three input datasets (RNA-seq, Proteomics, DNA methylation). Next, regulatory clusters were assigned using the cut-offs of adjusted p values < 0.05, and mRNA |*log*_2_*FC*| > 1.0, protein |*log*_2_*FC*| > 0.5 and |DNA methylation beta difference| > 0.1. The regulatory labels from grouping method 2 were chosen (**Table 1, Supplementary Table 2**). ORA was performed on each of the clusters using the background as defined by the background method (meaning all genes with p < 0.05 in at least two datasets). One point to note is that the state unchanged includes unmeasured; for example, a protein may be changed, but not detected, owing to technical limitations. Similarly, as we are using DNA methylation array data, a gene’s initial point of dysregulation may be at the methylation layer, yet not probed. Given the choice of filtering is specific to the biological question and technical setup, our package enables users to pre-filter and remove undetected genes at each layer.

### TF in each regulatory cluster

TFs were identified in each regulatory cluster using the TF database from DoRothEA^49^ (level A relationships). Relationships were retained only where the TF and the target had a significant (p < 0.05) change on the protein (TF) and mRNA (target) level, and the direction of change agreed with the direction of mode of regulation (MOR). For example, if the TF acts to enhance the gene target, the protein level of the TF should be positive and in turn the RNA level should also be positive. sciMoTF (version 0.1.1, https://github.com/ArianeMora/scimotf) was used to identify and visualise TFs that were enriched in clusters. Fisher’s Exact Test (FET) with Benjamini Hochberg (BH) correction was used, whilst the background consisted of all identified TFs in other regulatory clusters.

### ChIP HIF1A validation

ChIP-seq peaks from HIF1A (GSM3417826, GSM3417827, GSM2723878) and EPAS1 (GSM3417828, GSM3417829, GSE120887) were downloaded using ChipAtlas^72^ with a threshold of 50 (i.e. for peaks to be present they must have a Q value < 1E-05). These peaks were used to check whether HIF1A peaks did indeed occur at the TSS of the TPDE genes in patient derived renal cell carcinoma cell lines. IGV jupyter^73^ was used to view the peaks.

### Variational autoencoders to compress patient’s genes’ features

A variational autoencoder (VAE) is a generative machine learning method that can be used to learn to create a latent (i.e. compressed) representation of data. We choose a uniform gaussian as the prior distribution for the latent variables. The conditional distribution of the latent variables given the input data is given by the encoding *q*_*θ*_(*z*|*x*), and the conditional distribution of the data given the latent variables is represented by the decoding function *p*_*ϕ*_(*x*|*z*).Where the parameters, *θ*, and *ϕ*, include the weights and biases of the neural network that are learnt during training, via optimising with respect to an objective function. The objective function is used to 1) ensure the latent distribution approximates a standard gaussian by minimising the distance via mean maximum *D*_*MMD*_, 2) ensure the decoded data point is as close to the input as possible via mean squared error (MSE) i.e. we have a good generative model.

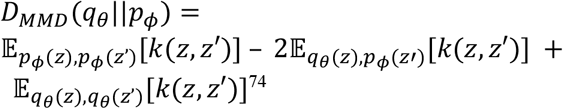

Above, the distance between the learnt distribution and the expected distribution is computed via the distance between distributions *q*_*θ*_ and *p*_*ϕ*_ using a kernel function, *k*. In our implementation we use a gaussian kernel, following the implementation from Zhao *et. al*^74^.

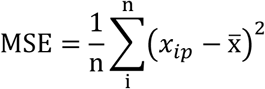

MSE is used to calculate the mean difference between the input data point, *x*_*ip*_, and the decoded value, *x*_*ip*_′. Where *x*_*ip*_is a set of features belonging to one patient, *p*, and one gene, *i*. Features selected as input were: protein levels, mRNA levels, protein, mRNA and methylation tumour versus normal difference, equating to a vector of seven features. Given the differing ranges of each feature, the data are scaled w.r.t gene *i* for a given data type prior to calculating the difference between tumour and normal. For example, we normalise *y*_*ip*_ as follows: *y* is the value for e.g. mRNA, *i* is the gene index, and p is the patient index, where min and max are calculated across all patients for gene *i*, omitting any missing data.

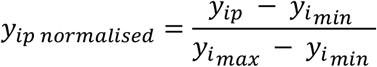

Prior to selecting the training set, genes with any feature with a z-score greater than two are considered outliers and omitted from the training set, however, are still input into the trained model. Our formulation lends to using the integrated value, *z*_*ip*_, as input to statistical functions such as Mann Whitney U, or a t-test. As an example, we show the formulation of the t-statistic on using integrated values, *z*_*ip*_, as defined in the results section. The test statistic for the difference between two patient groups, *S* and *G*, is thus defined as follows:

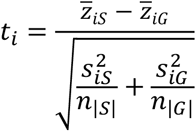

Significance is determined via two-sided null hypothesis, namely, that there is no difference between the means of the encoded values for patients belonging to group *S* and group *G*. In the non-parametric case, we use Mann Whitney U, where the null hypothesis that there is an equal chance that a patient from *S*, is greater than a patient from *G* and vice versa, as calculated by ranking each patient via their encoding.

### Creating the integrated dataset for the ccRCC case study

To identify genes that changed significantly across the three data types between patient groups we developed an integration package using a VAE (formalised above) to first integrate the data, reducing the features across patients to a single data point then perform statistics across two groups of patients. For the CPTAC dataset missing normal data were substituted with the mean normal value from patients with matching clinical information (stage and age group). Since methylation values weren’t matched, we use only the tumour methylation difference minus the average normal methylation from the TCGA patients. Only patients with matching protein and RNA normal and tumour data were used for training (59 cases). Data (genes by patients) from each SiRCle cluster were used to train a VAE (train-test split = 75:25), with 1 internal layer (5 nodes), and a single latent node, using a maximum mean discrepancy parameter of 0.25, mean squared error loss, rectified linear unit activation functions, Adam optimiser, batch size of 16, maximum epochs of 100 with early stopping after three epochs of no improvement in loss. These configurations were constant for all VAEs, and we end up with 10 trained VAEs corresponding to the 10 SiRCle clusters.

### Statistics on the integrated dataset

To determine genes that changed significantly between the groups of patients across the levels of information, we tested each gene using Wilcoxon rank sum test (given MDS_TMDE, TPDS_TMDE, MDE, MDE_TMDS, TMDS, TPDE_TMDS were not normally distributed) on three comparisons: 1) old (56) vs young (7), and 3) BAP1 (13) mutations vs PBRM1 (33) mutations, Stage 4 (10) vs Stage 1 (45) (**Supplementary Table 1, Supplementary Fig. 1a**). P-values were corrected using BH FDR on all genes within each cluster.

Before returning the values to the user the direction of the VAE integrated value with that of protein Log2FC, or RNA Log2FC (Supplementary **Fig. 5a**). To quantify the integrated change, termed “Integrated difference”, we computed the “distance” condition 1 (e.g. late stage) patients’ mean VAE value from condition 0 (e.g. early stage) mean VAE value (subtracting the value from condition 0 from condition 1 in an absolute sense). Note this is defined earlier.

We used a fGSEA (https://github.com/ctlab/fgsea) test to identify coordinated changes in metabolic signatures^50^, we use the integrated difference as the statistic provided to fGSEA and use 10,000 permutations.

### Single cell analysis

Gene counts, and cell annotations, and UMAP coordinates, were downloaded from Krishna *et al*.^43^. The single cell data were used to quantify the “co-expression” and “cell-types” of the genes changing between Stage IV and Stage I patients and BAP1 vs PBRM1 patient groups. We divide genes into the “positive” and “negative” groups by selecting genes significant (p < 0.05) and either a positive change (> 0) between the two populations or negative (<0).

We then test which of these genes are differentially expressed between the 31 cell types using the “scanpy.t1.rank_gene_groups” (as annotated by Krishna *et al*.^47^), one vs rest test, method=t-test. Using the returned p.adj and *log*_2_*FC* for each gene, we determined the genes associated with each cell type by selecting genes with a p.adj < 0.05 and a *log*_2_*FC* > 0.5 for that cell type. Finally, using the SiRCle integrated difference gene sets and the cell type gene groups, we tested for enrichment in each cell type for SiRCle Stage IV-Stage I and PBRM1-BAP1 set using a Fisher’s exact test, correcting p-values using BH FDR correction. The top 5 genes are presented in **Fig. 5a** as per odds ratio value. For heatmap and scatterplot visualisations genes are z-score normalised. Finally, using the Protein Atlas (https://www.proteinatlas.org/) we check the prognostic status of each of the genes and plot these using GEPIA2 (http://gepia2.cancer-pku.cn/) using the default quantile cut-offs (0.25, 0.75).

### Metabolomics profile comparing early and late stage

Median normalised metabolomic profiling data of 138 matched clear cell renal cell carcinoma (ccRCC) and normal tissue pairs was downloaded from the supplementary table 2 of Hakimi et al.^7^. Tumour samples were normalised to their matching normal tissue sample based on the matching index by calculating the fold change. Next, we performed differential metabolite analysis (DMA) comparing late stage (Stage 3 and 4) versus early stage (stage 1 and 2) and old (Age at surgery >58) versus young (Age at surgery <42) patients. Mann-Whitney U test (wilcox.test in R) was used to calculate the p-value and corrected using BH. Unnamed metabolites were removed, and metabolic pathways assigned using published pathway information^7^.

## Data availability

Data generated as part of this study are available as supplementary tables or via the Githu https://github.com/ArianeMora/SiRCle_multi-omics_integration, where information to download all raw data from the respective studies is available. If there are any additional information required for reanalysis of the data reported in this paper contact.

## Code availability

The SiRCle method described in this manuscript is available as a Python package on GitHub (https://github.com/ArianeMora/scircm), along with source code (https://github.com/ArianeMora/SiRCle_multiomics_integration). Installation instructions and documentation are available at https://arianemora.github.io/SiRCle_multiomics_integration/. If there are any additional information required for reanalysis or code adaptations, please post an issue on the GitHub.

## Acknowledgements

We thank Connor Rogerson, Lorea Valcarcel-Jimenez, Dylan Ryan, Marco Sciacovelli, Ming Yang, Sanjana Tule, Sam Davis, Oliver Hughes, and all other members of the Frezza and Bodén lab for discussion and literature advice. The results presented in this study are based, in part, on data generated by The Cancer Genome Atlas (TCGA), the TCGA Research Network, the Clinical Proteome Tumor Analysis Consortium (CPTAC), the study of Clark et al. (Cell 2019), Hakimi et al. (Cancer Cell 2016) and Krishna et.al. (Cancer Cell 2021). We thank them for making their data publicly available.

C.S work was funded by the European Union’s Horizon 2020 research and innovation programme under the Marie Skłodowska-Curie grant agreement No 722605., the CRUK Programme Foundation award (C51061/A27453) and the Alexander von Humboldt Professorship to C.F. C.F was supported by the Medical Research Council (MRC_MC_UU_12022/6), the CRUK Programme Foundation award (C51061/A27453), ERC Consolidator Grant (ONCOFUM, ERC819920) and by the Alexander von Humboldt Foundation in the framework of the Alexander von Humboldt Professorship endowed by the Federal Ministry of Education and Research. A.M. and B.B. were funded by Australian Government Research Training Program.

## Contributions

C.S., A.M. and M.B. designed the SiRCle method. A.M. and C.S. implemented code. C.S, C.F. and A.M. interpreted the biological readouts. B.B analysed the single cell data. A.M. and C.S. wrote the manuscript and prepared the figures. C.F., M.B. and B.B. reviewed and edited the manuscript. C.F. and M.B lead the investigation.

## Supplementary information

### Supplementary Figures

**Supplementary Fig. 1.**
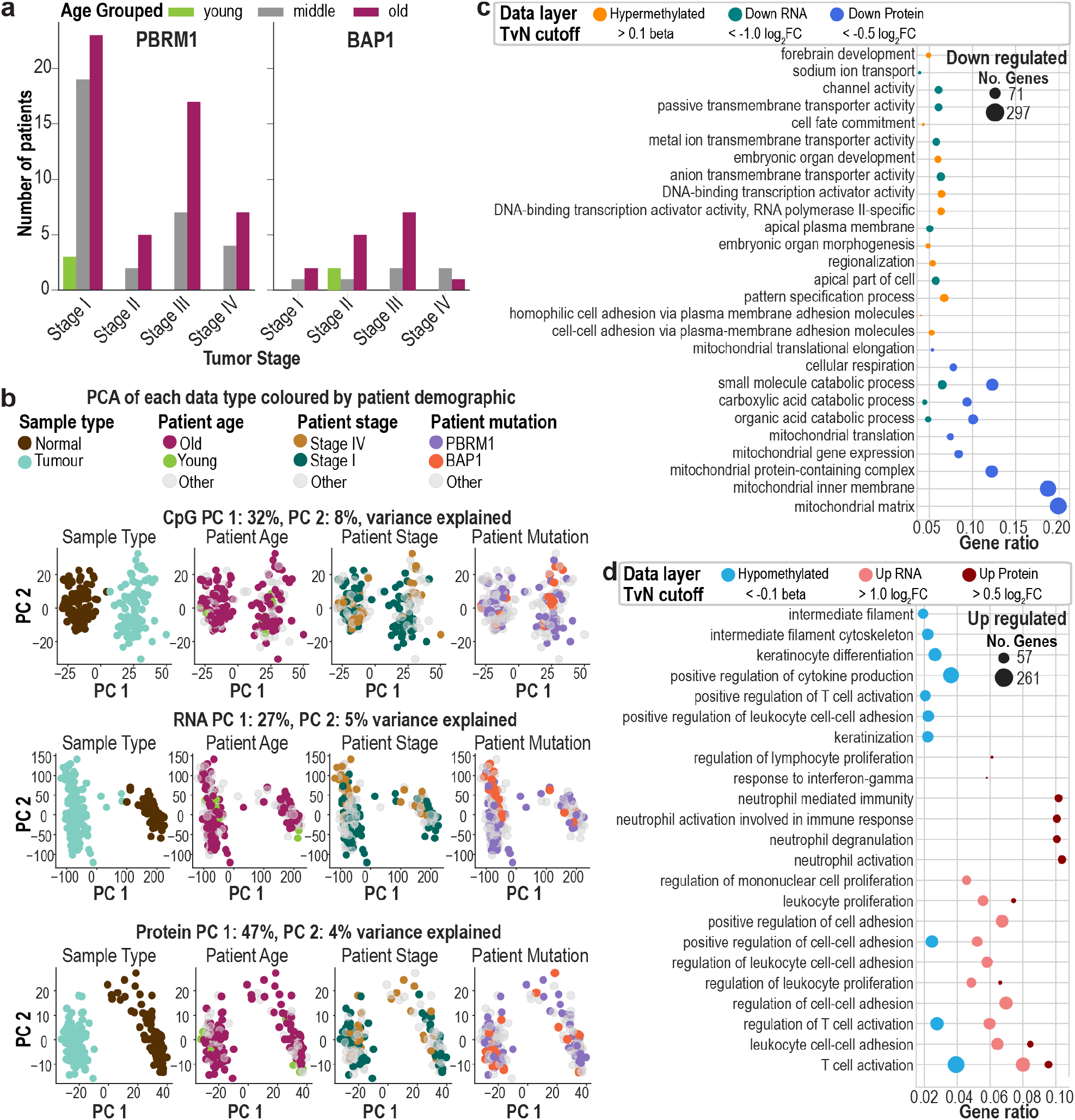
**a**, Case annotations based on mutation, tumour stage and patients age. **b**, Principal Component Analysis (PCA) shows the largest source of variation in all datasets across patient samples is the Sample Type while patient annotations don’t account for the top two principal components. Note where an annotation is defined on the tumour level, for example stage or mutation, that annotation is also used to colour the normal sample from the same patient. **c**, The top 10 gene ontology terms from over representation analysis for each data layer are visualised. Where two dots appear on the same line, it means they were both top terms for that data layer. This only occurs for the up regulated terms, i.e. the terms that were associated with an increase on the protein or mRNA layers. Data layer refers to the data layer on which the differential analysis was run, and TvN cut-off denotes the cut-off used when selecting genes for ORA. **d**, as in c, except with the bottom terms.

**Supplementary Fig. 2.**
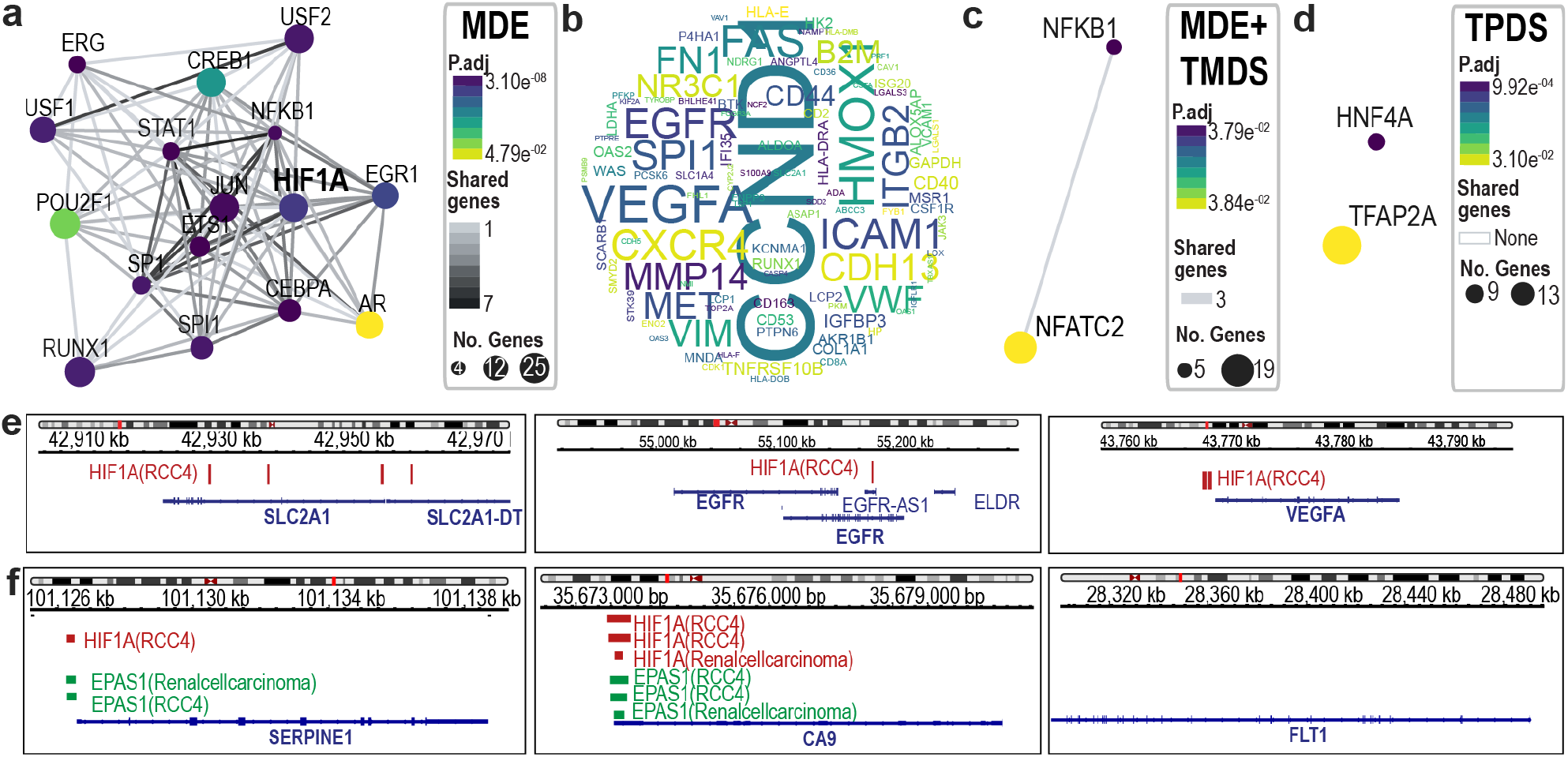
**a**,**c-d** Transcription factor (TF) network of TFs from manually curated repositories (DoRothEA) that may drive genes in the cluster Methylation Driven Enhancement (MDE) **(a)**, MDE+ Translation and post-translational Modification Driven Suppression (TMDS) **(c)** and Transcription and Processing Driven Suppression (TPDS) **(d)**. The dot size corresponds to the number of genes of the cluster that are targeted by the TF. The colour of the dot shows the p-adjusted value (p.adj). The connecting lines (grey) shows the number of common genes the connected TFs regulates. **b**, Wordcloud including the TF targets with shared TF binding in the MDE cluster, with the size corresponding to how many different TF’s are predicted to regulate the genes expression. **e**, HIF1A binding for genes in RCC cell in the MDE cluster. **f**, HIF1A and EPAS1/HIF2A binding for genes in the TPDE cluster.

**Supplementary Fig. 3.**
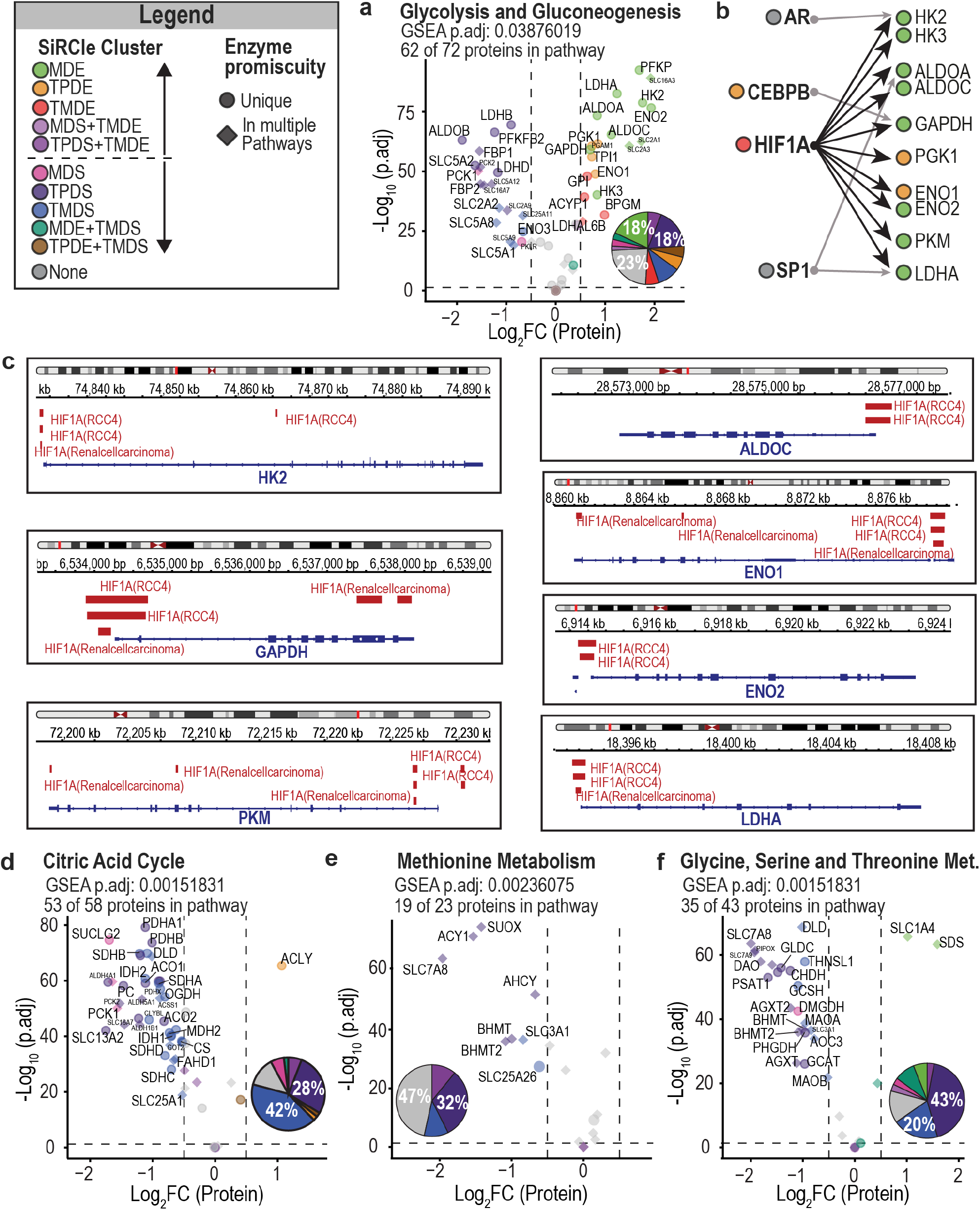
Volcano plots are based on the metabolic signatures and the number of proteins detected of the pathway are reported. Proteins that are unique for the metabolic pathways are displayed in circles and proteins that are part of multiple metabolic pathways are displayed in diamond. The colour code depends on the SiRCle cluster the protein is part of and is summarised in the pie chart. The protein Log2FC is calculated between tumour versus normal. **a**, Volcano plot of glycolysis and gluconeogenesis. **b**, Transcription factor (TF) factor network of TFs from manually curated repositories (DoRothEA^49^) that drive glycolytic enzymes in the cluster Methylation Driven Enhancement, MDE and Transcription and Processing Driven Enhancement, TPDE. **c**, HIF1A ChIP-seq peaks binding sites at glycolytic enzyme TSS’s. **d**, Volcano plot of the citric acid cycle. **e**, Volcano plot of methionine metabolism. **f**, Volcano plot of glycine, serine and threonine metabolism.

**Supplementary Fig. 4.**
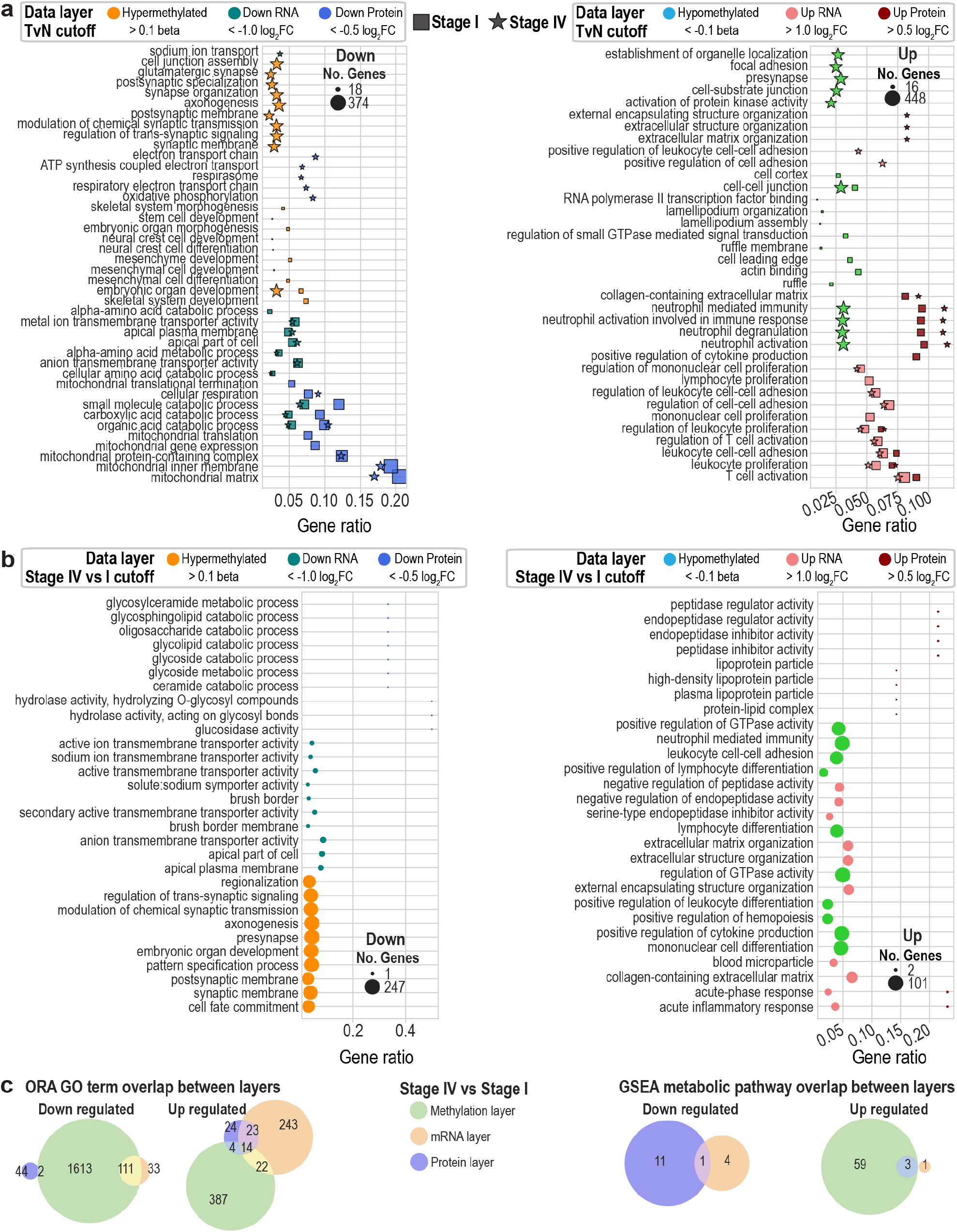
**a**, Top 10 gene ontology (GO) terms from over representation analysis (ORA) for each data layer are visualised for stage I tumour vs normal and stage IV tumour versus normal (TvN) on the same plot. Where two dots appear on the same line, it means they were both top terms for that data layer or in both stage I and stage IV TvN comparisons. Data layer refers to the data layer on which the differential analysis was run and the state, for example increased or decreased in tumour (Up and Down respectively), and TvN cut-off denotes the TvN cut-off used when selecting genes for ORA. **b**, Top 10 GO terms from ORA for each data layer are visualised for stage IV tumour vs stage I tumour. **c**, overlap between all significantly enriched GO terms and metabolic pathways using gene set enrichment analysis (GSEA) with the test statistic from the differential analysis for a given layer used as input. An overlap indicates that a pathway or term was significantly changed in both data layers. “Up regulated” refers to a positive enrichment score whilst “Down regulated” refers to a negative enrichment score for protein and mRNA layers, whilst the opposite for the DNA methylation layer.

**Supplementary Fig. 5.**
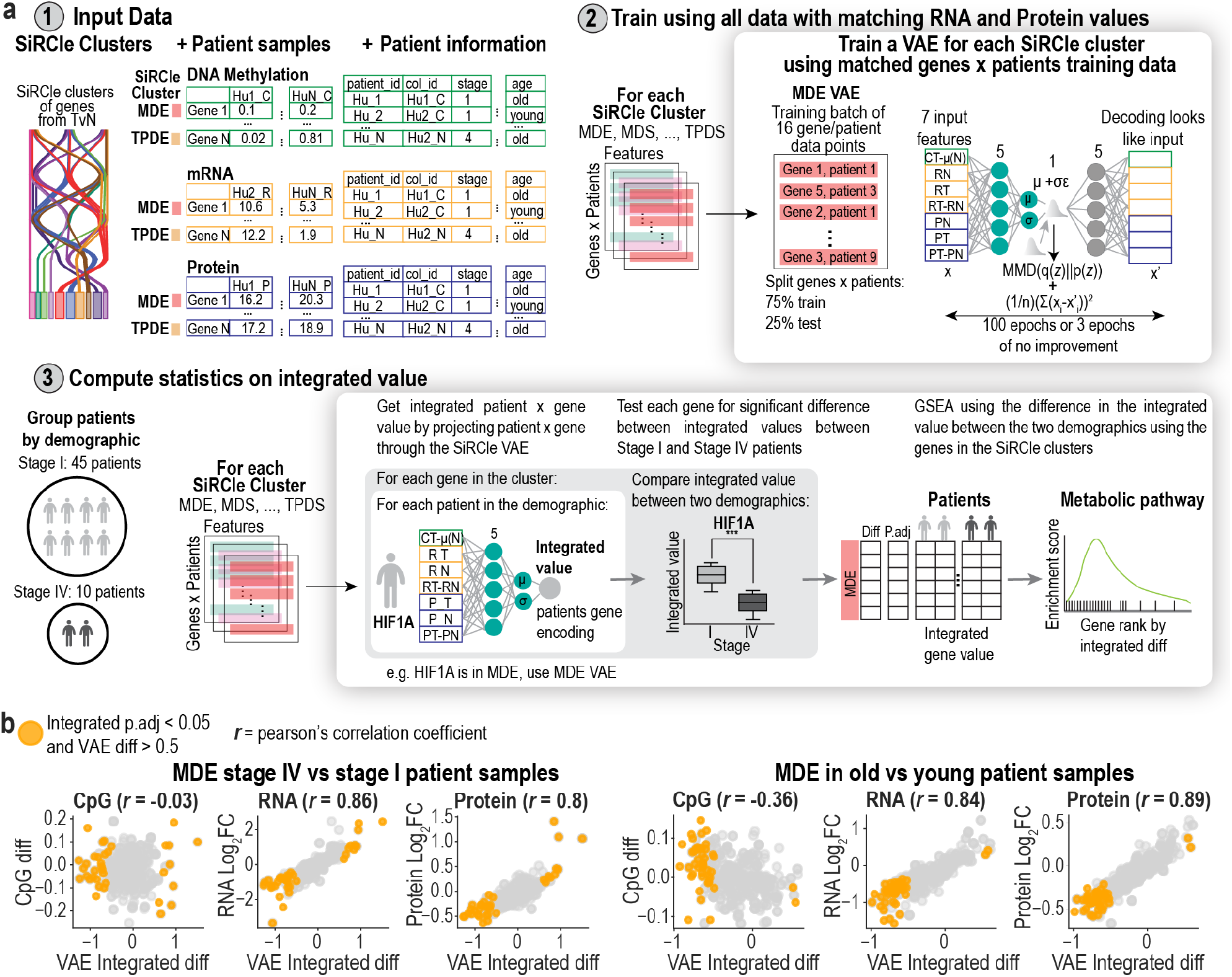
**a**, Overview of SiRCle integration using a variational autoencoder (VAE) and statistics on the VAE dimension. **b**, Dispersion between biological features and the integrated difference in the Methylation Driven Enhancement (MDE) SiRCle cluster. The integrated difference is the difference between patients of two groups, similarly, the integrated p.adj is the adjusted p-value from the integrated difference between two patient groups (e.g. stage IV vs stage I). Each point represents a gene, and the difference is the mean difference between the groups.

**Supplementary Fig. 6.**
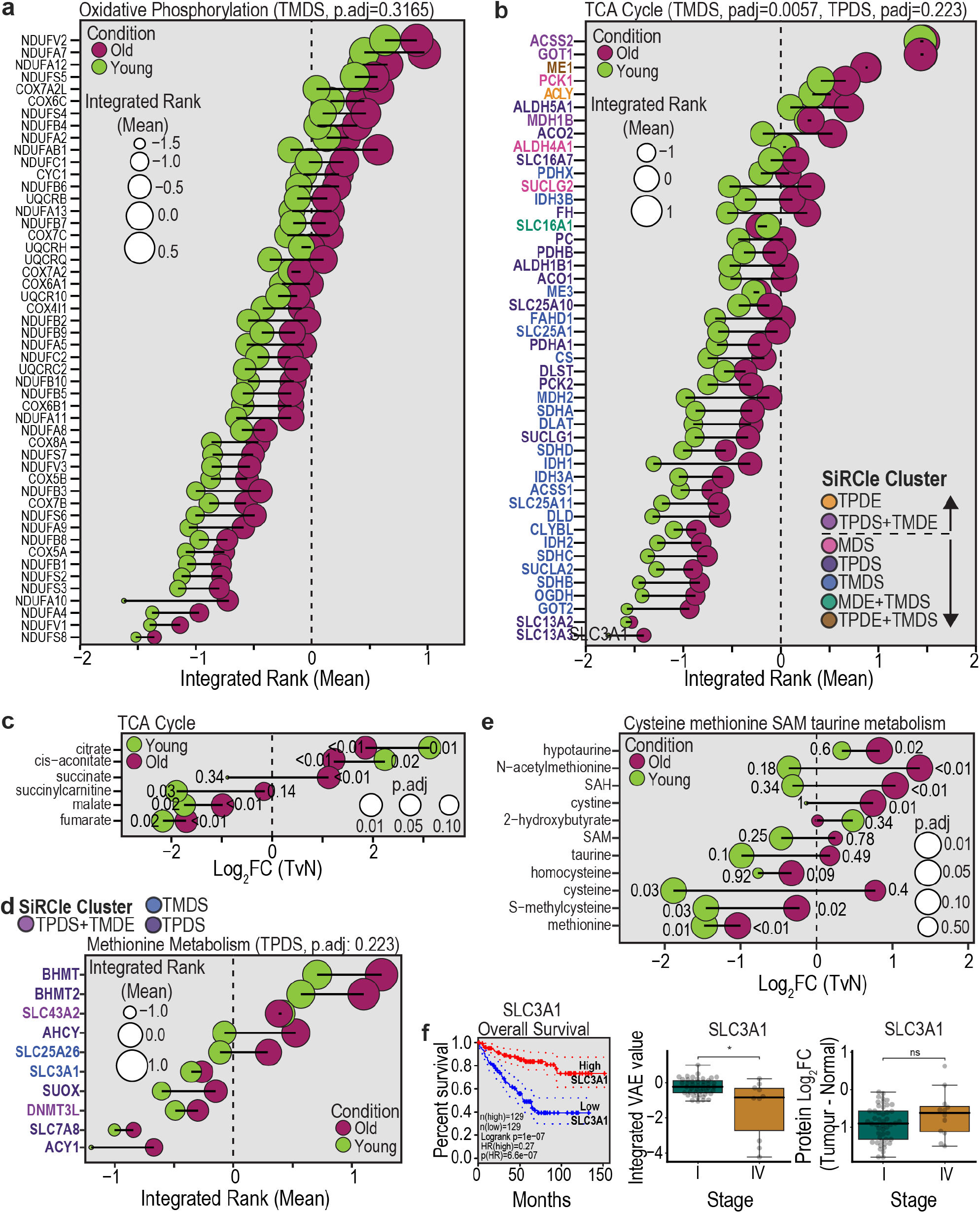
Metabolic signatures based on gene expression are defined by Gaude *et. al*^50^ with p.adj values corresponding to the GSEA results after ranking the genes in each SiRCle cluster using the VAE integrated rank. Metabolite pathways based on metabolites are defined by Hakimi *et. al*^7^. **a**, Comparison of the VAE integrated rank of old with young patients for oxidative phosphorylation genes that are part of the Translation and post-transcriptional Modification Driven Suppression (TMDS) SiRCle cluster. **b**, Comparison of the VAE integrated rank of old with young patients for TCA cycle genes colour coded for the SiRCle clusters they are part of. **c**, Comparison of the log_2_FC tumour versus normal (TvN) of old with young patients for TCA cycle metabolites. **d**, Comparison of the VAE integrated rank of old with young for genes corresponding to the methionine metabolism pathway; colour coded for the SiRCle clusters they are part of. **e**, Comparison of the log_2_FC TvN of old with young patients for “cysteine, methionine SAM, taurine metabolism” metabolites. **f**, *SLC3A1* survival curve based on (http://gepia2.cancer-pku.cn/) using quantile default cut offs of 25%, 0.75%. Integrated difference and protein log_2_FC difference are comparing stage I with stage IV patients.

**Supplementary Fig. 7.**
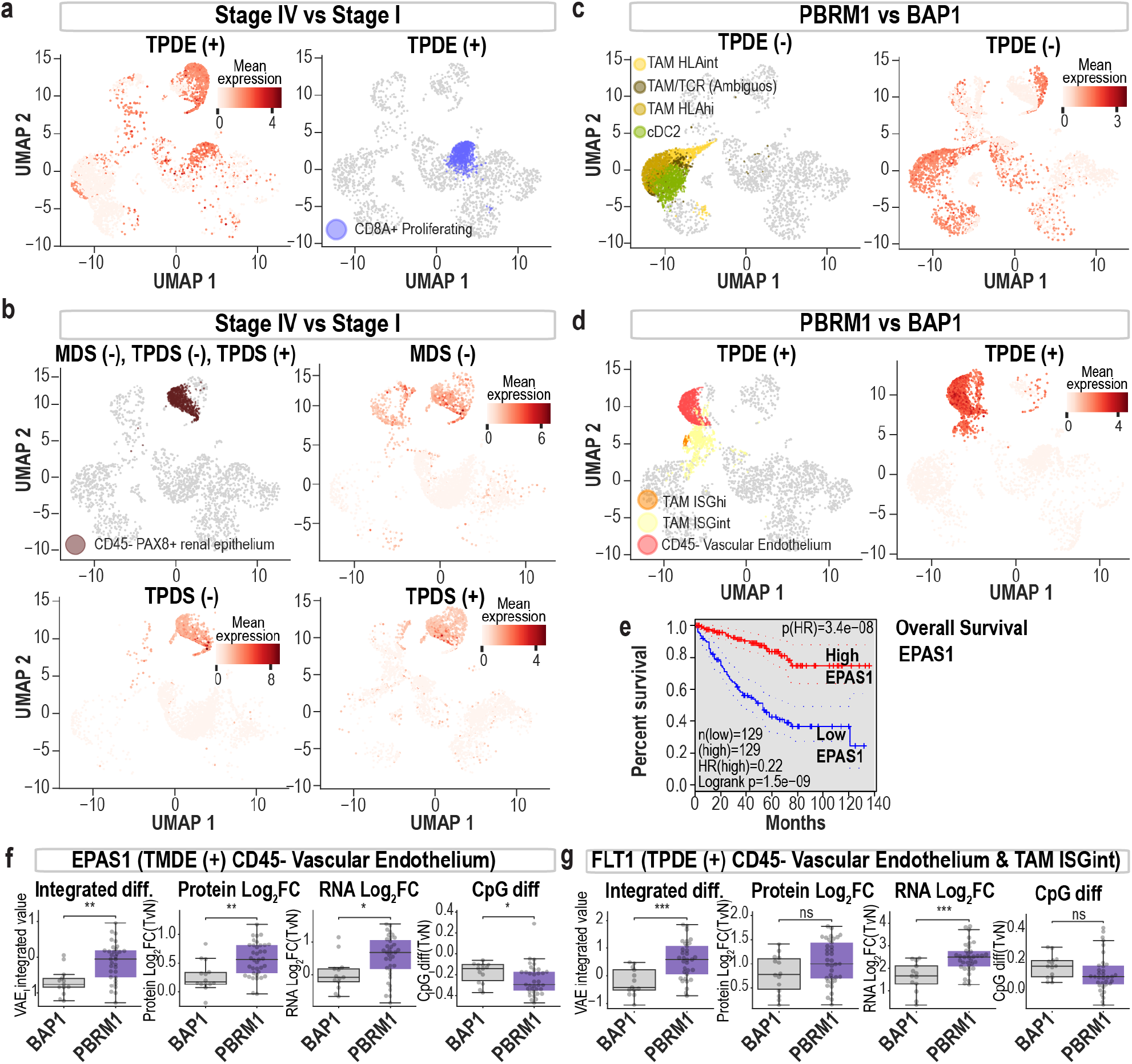
UMAP plots use the UMAP cell coordinates and cell annotations from Krishna *et. al*^47^ .While the gene expression is the mean gene expression for the genes identified as enriched in any cell type and the given SiRCle cluster subset into (+) for significant (p <0.05) changes with stage IV > stage I, PBRM1 > BAP1, and (-) stage IV < stage I, PBRM1 < BAP1. Boxplots show the integrated VAE value for each patient with either a BAP1 or PBRM1 mutation, alongside the protein and RNA Log2FC, and methylation beta difference between tumour and normal for that same group of patients. Boxplot titles include the SiRCle cluster and the cell type(s) it is enriched in. Survival plots were plotted using GEPIA2 (http://gepia2.cancer-pku.cn/) using quantile default cut-offs of 25%, 0.75%. **a**, UMAP plots for TPDE (+). **b**, UMAP plots for stage IV vs stage I, Methylation Driven Suppression, MDS (-), Transcription and Processing Driven Suppression, TPDS (-) and TPDS (+), enriched in CD45-PAX8+ renal epithelium. **c**, UMAP plots for PBRM1 vs BAP1 Transcription and Processing Driven Enhancement, TPDE (-). **d**, UMAP plots for PBRM1 vs BAP1 Translation and post-transcriptional Modification Driven Enhancement, TMDE (+). **e**, Survival EPAS1/HIF2A. **f**, *EPAS1*/*HIF2A* VAE integrated difference alongside tumour vs normal summaries for patients with either *BAP1* or *PBRM1* mutations. **g**, *FLT1* VAE integrated difference alongside tumour vs normal summaries for patients with either BAP1 or PBRM1 mutations.

## Supplementary Tables

**Table 1: Clinical information of patients from each study**

- Sheet 1: Summary of CPTAC and TCGA patients and their demographics
- Sheet 2: Sample IDs removed based on QC

**Table 2: DE results for each dataset**

- Sheet 1: Protein DA for TvN
- Sheet 2: RNA DE for TvN
- Sheet 3: DNA methylation DCpG for TvN
- Sheet 4: Protein enrichment (ORA for UP genes)
- Sheet 5: Protein enrichment (ORA for DOWN genes)
- Sheet 6: RNA enrichment (ORA for UP genes)
- Sheet 7: RNA enrichment (ORA for DOWN genes)
- Sheet 8: CpG enrichment (ORA for UP genes)
- Sheet 9: CpG enrichment (ORA for DOWN genes)

**Table 3: Results from the SiRCle clusters**

- Sheet 1: RCM output, which includes the mRNA, protein, & DNA methylation Log2FC’s along with the cluster assigned.
- Sheet 2: ORA outputs for each cluster

**Table 4: TF analysis output**

- Sheet 1: Transcription factor output table

**Table 5: Metabolomics data for tumour vs normal**

- Sheet 1: clinical information for metabolomics patients
- Sheet 2: TvN differential expression results
- Sheet 3: GSEA on protein TvN for metabolic pathways
- Sheet 1: comparison of Stage IV vs Stage I
- Sheet 2: comparison of young vs old

**Table 6: ORA and GSEA on Stage I and Stage IV samples**

- Sheet 1: Protein DA for Stage I TvN
- Sheet 2: RNA DE for Stage I TvN
- Sheet 3: DCpG for Stage I TvN
- Sheet 4: DE analysis for Stage IV TvN
- Sheet 5: DA analysis for Stage IV TvN
- Sheet 6: DCpG analysis for Stage IV TvN
- Sheet 7: DE analysis for Stage IV Tumour v Stage I Tumour
- Sheet 8: DA analysis for Stage IV Tumour v Stage I Tumour
- Sheet 9: DCpG analysis for Stage IV Tumour v Stage I Tumour
- Sheet 10: ORA UP for TvN Stage I and Stage IV
- Sheet 11: ORA DOWN for TvN Stage I and Stage IV
- Sheet 12: ORA UP for Stage IV v Stage I tumour
- Sheet 13: ORA DOWN for Stage IV v Stage I tumour
- Sheet 14: GSEA on metabolic pathways Stage IV v Stage I tumour Positive NES
- Sheet 15: GSEA on metabolic pathways Stage IV v Stage I tumour Negative NES

**Table 7: Integrated dataset for patients**

- Sheet 1: genes x patients with integrated value
- Sheet 2: comparison of Stage IV vs Stage I
- Sheet 3: comparison of young vs old
- Sheet 4: comparison of BAP1 vs PBRM1

**Table 8: GSEA results for comparisons**

- Sheet 1: GSEA on metabolic pathways Stage IV vs Stage I
- Sheet 2: GSEA on metabolic pathways of young vs old
- Sheet 3: GSEA on metabolic pathways of BAP1 vs PBRM1

**Table 9: Single cell output of enriched groups**

- Sheet 1: comparison of Stage IV vs Stage I
- Sheet 2: comparison of BAP1 vs PBRM1

